# The Microbiome of Bioreactors Containing Mass-Cultivated Marine Diatoms for Industrial Carbon Capture and Utilization

**DOI:** 10.1101/2023.11.01.565100

**Authors:** Nerea Johanna Aalto, Ingeborg Hulda Giæver, Gunilla Kristina Eriksen, Linn Israelsen, Stina Krsmanovic, Sebastian Petters, Hans C. Bernstein

**Author notes:** Correspondence: Hans C. Bernstein, UiT-The Arctic University of Norway, BFE-fak., Postboks 6050 Langnes, 9037 Tromsø, NORWAY.

## Abstract

Marine microalgae are a promising innovation platform for carbon capture and utilization (CCU) biotechnologies to mitigate industrial greenhouse gas emissions. However, industrial-scale cultivation of algal mono-cultures is challenging and often unscalable. Non-axenic microalgae in large semi-open photobioreactors lead to the co-cultivation of diverse microbial communities. There is limited knowledge about the “bioreactor ecology” involving microalgae interacting with the microbiome and its subsequent impact on process stability and productivity. In this study, we describe the semi-continuous industrial mass cultivation of the cold-adapted marine diatom, *Porosira glacialis* UiT201, by investigating the prokaryotic and microeukaryotic (phytoplankton and heterotrophic protist) communities. Data were collected in two consecutive time series experiments, representing the initiation and operation of an preindustrial scale CCU photobioreactor (300,000 liters). The first experiment experienced a culture “crash” of the focal strain after 39 days, while the second culture remained stable and “healthy” for 60 days. The results highlight that this mass cultivation system represents a unique industrial marine microbial ecosystem. The succession of the prokaryotic community was primarily driven by species replacement, indicating turnover due to selective bioreactor conditions and/or biological interactions. Nonetheless, the bioreactor consistently harbors a recurring and abundant core microbiome, suggesting that the closely associated bacterial community is influenced by microalgae-specific properties and can endure a dynamic and variable environment. The observed culture collapse of *P. glacialis* coincided with changes in the core microbiome structure and different environmental growth conditions compared to the stable and “healthy” experiment. These findings imply that cohabiting microbial taxa within industrial microalgae cultivation likely play a critical role in stabilizing the conversion of industrial CO_2_ into marine biomass, and changes in community structure serve as an indicator of process stability.

## 1 Introduction

Growing levels of atmospheric carbon have an impact on biodiversity, human health, the economy, and the global geopolitical landscape. As the world’s climate nears new tipping points [1], it is imperative that we not only reduce emissions but also take new steps toward carbon capture and utilization (CCU). Innovative biotechnologies that capitalize on photosynthetic carbon fixation, which naturally controls global carbon cycles, will contribute significantly to the solution [2]. Marine microalgae are specifically promising because they have a much higher carbon fixation capacity than plants. Living microalgae make up just 1-2% of global biomass, but they account for almost 50% of the world’s primary production [3, 4]. The utility of microalgae for CCU is being demonstrated at an industrial installation in (Arctic) northern Norway, that harnesses cold-adapted marine diatoms to capture CO_2_ at the source from an operational ferrosilicon smelter plant [5]. This is a prominent example of how innovative algal biotechnology can be deployed for CCU by taking advantage of locally isolated diatom strains adapted to the seasonality and seawater resources available to coastal industry locations. A recent study reported a difference in the photophysiology of the Arctic diatom *Porosira glacialis* UiT201 compared to temperate microalgae that enables strong photosynthetic flexibility in the dynamic light and temperature environment, as is uniquely experienced in coastal, Arctic regions [6].

Scalingup mass cultivation of microalgae for CCU must, however, overcome several barriers that include managing the microbial populations within large industrial scale photobioreactors. As axenic mass cultivation of microalgae is technically challenging, large photobioreactors are often populated by multi-domain microbial communities and function as their own unique ecosystems [7, 8]. A better understanding of how these communities assemble and behave under stable and unstable – i.e., reactor crash – conditions is needed to scale bioreactors up to volumes capable of effective industrial CO_2_ capture. Open airlift column bioreactors are effective for scaled-up industrial operation, land space utilization, and improved CO_2_ gas-to-liquid transfer [5]. However, the challenge linked to open-systems is a constant exposure to the environment and thereby capacity of recruiting new microbial taxa to the system [8]. The extent to which a bioreactor recruits microbial taxa from the environment depends on how the inputs (e.g., water, nutrients, innocula) and control volume are exposed to the environment. This presents a unique intersection between bioprocess engineering for CCU and microbial ecology.

Coinhabiting microbial taxa within industrial photobioreactors are not necessarly harmful. Infact, previous studies have demonstrated that the majority of the interspecific interactions between microalgae and bacteria are mutualistic [9–11]. The beneficial associations are based on exchange of synthesized compounds enhancing growth of both partners [12, 13]. The invasion of other microbial eukaryotes i.e., phytoplankton and heterotrophic protist (hereafter referred as “microeukaryotes”) instead can cause a rapid culture collapse of the focal strain [8]. Some mass cultivation attempts used euryhaline and high pH tolerate focal strain to prevent the contamination of competitive or predative species [14–16]. However, extreme conditions reduce the number of potential strains and might complicate the operational processes. To uncover the potential and risks associated with the coexisting microbiome, a shift in focus from focal strain-centered investigations to ones that encompass the entire biological component of the production system is necessary [17].

Major knowledge gaps exist in our understanding of how to control or even measure the complex interactions between microalgae and bacteria. Only few previous studies have measured the microbial diversity of photobioreactors and none (to our knolwege) have investigated the effects on the stability of CCU-specific operations. In order to address some of these knowledge gaps, mass cultivation of the marine centric diatom *Porosira glacialis* UiT201 in a preindustrial scale CCU operation was combined with an investigation of the community composition and temporal dynamics of the bioreactor microbiome. This study systematically explores three research questions: (i) Which prokaryotic and microeukaryotic taxa are recurrently observed in mass cultivation alongside the focal diatom strain? (ii) What are the discernible succession patterns within the prokaryotic and microeukaryotic consortia of the bioreactor microbiome, and how are these patterns influenced by the bioreactor’s fluctuating environmental conditions? and (iii) In what ways do the outcomes of two sequential time-series experiments, one concluding with a premature culture crash at 39 days and the other achieving sustained culture stability for 60 days, converge or diverge in terms of microbiome structure and bioreactor growth conditions?

## 2 Materials and Methods

### 2.1 Algae inoculum

The focal strain used to inoculate bioreactors was the centric diatom *Porosira glacialis* (UiT-201). It was originally single-cell isolated from a sediment sample collected in the Barents Sea (N 76° 27.54′, E 033° 03.54′) in 2014 [18]. The monoculture was maintained in a temperature regulated incubator (Termaks KBP 6151) at 8 °C with a scalar irradiance of 65 µmol m^-2^ s^-1^ and diluted weekly with filtered (0.45 µm) and autoclaved seawater containing inorganic marco- and micronutrients from a plant nutrient mix YaraTera Kristalon Purple (Yara, Norway) and sodium metasilicate pentahydrate (Na_2_SiO_3_×5H_2_O; Permakem, Norway).

### 2.2 Photobioreactor operation

Mass cultivation of *P. glacialis* took place in northern Norway (69.22°N; 18.08°E) in two consecutive time series experiments: 19.08-27.09 and 19.10-17.12 in 2021. Hereafter the first time series recording is referred to as “T1-crash” due to unplanned discontinuation of cultivation caused by contamination and collapse of *P. glacialis* culture and the second as “T2-healthy” in which the focal strain was maintained at targeted abundance and growth rates until the end of time series. The cultivation platform is integrated into an industrial ferrosilicon smelting plant, Finnfjord AS, in collaboration with UiT – The Arctic University of Norway. Detailed description of the mass cultivation design and operation is given in Eilertsen et al. [5]. The platform contains four vertical column airlift photobioreactors with volumes of 6 m^3^, 6 m^3^, 14 m^3^, and 300 m^3^ [5]. This study deployed the largest 300 m^3^ reactor operated with culture volume of 150 m^3^ under semicontinuous conditions. Seawater for cultivation medium was obtained from an adjacent fjord at the depth of 25 m and filtered through a multichannel filtration unit with 0.45 µm as a smallest cartridge pore size and through a UV-sterilization unit (Model: MR3-350PP; Ultraaqua, Denmark). The sea water inflow to the reactor is however not sterile, despite the robust filtration system. Compressed air and flue gas from the factory with CO_2_ content of 3-7% were added into the bottom of the reactor using a rotating gas dispersing device [5]. Flue gas was sparged continuously during the cultivation periods except for three dates: 28.10., 08.12. and 14.12.2021. Illumination was provided for the reactors by ambient sunlight plus submerged LED lights. Additional artificial illumination was provided by a collection of 200 x 50W LED light units, placed above the culture surface of the bioreactor. The average light intensity in the 300 m^3^ reactor, provided by the collection of sources, was 36 µmol m^-2^ s^-1^. Inorganic nutrients (nitrite, nitrate and phosphate) a plant nutrient mix YaraTera Kristalon Purple (Yara, Norway), and Na_2_SiO_3_×5H_2_O (Permakem, Norway) were added manually to the culture from the start of T1-crash and following 9-days (19.10.-29.10.2021). Thereafter, the nutrients were automatically added using a dosing pump to ensure continuous addition of nutrients. On average, 1231 g 24 h^-1^ of Kristalon Purple and 1321 g 24 h^-1^ of Na_2_SiO_3_×5H_2_O were dosed as liquid solutions into the bioreactor.

### 2.3 Inoculation of the 300 m^3^ photobioreactor

Prior to each time series experiment, the inoculation volume of *P. glaciali*s was increased in two smaller (6 m^3^ and 14 m^3^) bioreactors. The 300 m^3^ bioreactor was pre-filled with ∼45 000 L of filtered seawater before the inoculum was introduced. Thereafter, the volume was increased for the next two days until a volume of 150 m^3^ was reached as this was the maximum operation capacity for the light delivery system. Semicontinuous operation was maintained by continuously adding filtered seawater for five days a week (Monday – Friday morning) with a mean inflow of 2100 L h^-1^. The culture surplus was extracted from overflow (150 m^3^) and algae was harvested.

### 2.4 Sample collection and growth condition monitoring

Daily measurements were obtained during scheduled work shifts (Monday-Friday) from the 300 m^3^ bioreactor for: temperature (C°; Endress-Hauser sensor iTHERMA ModuLine TM131 w) and dissolved oxygen (DO mg L^-1^; WTWMulti 360 meter with CellOx 325 sensor, Xylem Anlaytics, Welheim, Germany) from subsurface at ∼1 m depth. Liquid samples for cell counts were collected daily and for analysis of inorganic nutrients every second day. Samples were collected twice weekly (Monday and Friday) for genomic DNA isolation and dissolved organic carbon (DOC mg L^-1^). These samples were also collected from the 300 m^3^ bioreactor pre-filled with filtered intake water (hereafter referred as “SW bioreactor”), from inoculum (sample from 6 m^3^ and 14 m^3^ were combined), and once or twice from intake water used to semicontinuously dilute the reactor.

### 2.5 Sample processing and analyses

Concentration of inorganic nutrients of nitrite-nitrate (NO_2_^-^+NO_3_^-^), phosphate (PO_4_^3-^), and silicate (Si(OH)_4_) were analyzed immediately after filtering sample through a GF/C microfiber filter (Whatman). Nutrient specific Spectroquant kit (Merck Millipore, US) was used and the concentration was determined spectrophotometrically using a 96-well plate with a microplate reader (FilterMax F5, Molecular Devices, CA) in which the absorbance was measured at 540 nm. DOC samples were filtered through 0.2 µm cellulose syringe filters (Corning®) into acid washed and pre-combusted 60 mL glass vials and stored in dark at +4 °C for 12 months before acidification with 250 µl of 2N HCl to determine the concentration with TOC-VPH analyzer (Shimadzu, Japan). Modified Utermöhl method was used to obtain an estimation of *P. glacialis* cell density. Cells from two separate transects were counted from a 4-well dish with inverted light microscope (Zeiss PrimoVert) after 2 h of settling (Nunc, Thermo Fisher Scientific, US). Depending on the culture density up to 1 L of sample was filtered through each (n = 5 per sampling day) 0.22 µm Sterivex polyethersulfone membrane filter unit (Millipore) to obtain biomass for DNA extraction and following amplicon sequencing of 16S and 18S small subunit ribosomal RNA (SSU rRNA) gene. The filters were immediately stored at −80 °C for 3 months before DNA extraction took place at the UiT – The Arctic University of Norway.

### 2.6 Microbiome – DNA extraction and Illumina amplicon sequencing

DNeasy Power Water Kit (Qiagen, Carlsbad, CA) was used for the purification of genomic DNA. DNA was extracted from the collected biomass following the manufacturer’s protocol and stored at −80 °C until PCR. DNA quality and quantity were assessed by the NanoDrop™ Spectrophotometer and Qubit fluorometer (Invitrogen TM). Bacterial communities were identified following the Earth Microbiome Project (EMP) [19] 16S Standard Illumina library preparation protocol, using the forward-barcoded 515f (5′-3′ GTGYCAGCMGCCGCGGTAA) [20] and 806r (5′-3′ GGACTACNVGGGTWTCTAAT) [21] primers targeting the V4 region of 16S SSU rRNA. Targeted gene sequencing of microeukaryotes was determined following the EMP 18S Illumina Amplicon Protocol using the universal Euk_1391f (5′-3′ AATGATACGGCGACCACCGAGATCTACAC) and the barcoded EukBr (5′-3′ TGATCCTTCTGCAGGTTCACCTAC) primers targeting the V9 region of the 18S SSU rRNA [22, 23]. The PCR products were confirmed by gel electrophoresis in 1% agarose gels and visualized with GelRed® staining. Sequencing was performed on an Illumina MiSeq instrument (Illumina, San Diego, CA) at the Environmental Sample Preparation and Sequencing Facility at Argonne National Laboratory (Lemont, IL, USA).

### 2.7 Microbiome – Bioinformatic pipeline

Forward and reverse Illumina reads were imported into QIIME2 and thereafter demultiplexed using the complementary barcode file, for each sequencing run, respectively. These steps were executed using the EMP paired-end flag. DADA2 V2021.0 algorithm within QIIME2 was used for denoising and merging the reads [24]. During the denoising steps of the DADA2 package, the reads were pooled, filtered and chimera-checked. Additionally, the replicated reads were discarded. After removing the chimeric sequences and singletons, the reads of the 16S runs and 18S runs were merged, respectively. The DADA2 statistic on sequence reads is provided in Supplementary Data S1. Subsequently, the DADA2 package was used for inferring and filtering amplicon sequence variants (ASVs). Taxonomy was assigned by aligning these ASV sequences against the SILVA V138.1 database, using a 16S and 18S rRNA self-trained classifier [25, 26]. Classifiers were trained using RESCRIPt [27]. Although the 16S and 18S based taxonomic classification within SILVA V138.1 database is a standardized approach, it has limitations regarding classification and taxonomic resolution. It is noted that manual correction of taxonomic annotation was beyond the scope of this study. Nevertheless, order *Enterobacterales* was manually changed to order *Alteromonadales* as the main classified families obtained within *Enterobacterales* in our 16S dataset were *Colwelliaceae*, *Alteromonadaceae* and *Pseudoalteromonadaea* that are known to be members of *Alteromonadales* [28]. The phylogenetic rooted tree, generated with QIIME2 using MAFFT algorithm, was used for downstream diversity analysis.

### 2.8 Microbiome – Diversity analysis

All downstream analyses were completed using R [29]. The microbiome time-series data was analyzed using the “microeco” package (v16.0) [30] unless another package is specifically mentioned. Prior to community analysis, all reads that belonged to mitochondria, chloroplast and kingdom *Eukaryota* or were unassigned (no assigned kingdom) were removed from the 16S dataset. Filtering from the 18S dataset removed ASVs assigned to the kingdoms of *Bacteria* and *Archae*a and phyla of *Vertebrata*, *Arthropoda*, *Nematozoa*, *Annelida*, *Cnidaria*, *Mollusca*, *Tunicate*, *Phragmoplastophyt*a and *Basidiomycota* as well as unassigned reads. Both datasets, 16S and 18S, contained a high number of ASVs with low number of reads. For example, 2074 ASVs found just in 300 m^3^ bioreactor comprised only 0.6% of the total reads within 16S dataset. As we are only considering taxa with higher abundance, an abundance-filter was used to remove rare and possibly ambiguous ASVs from the datasets. The filter-threshold included only ASVs which mean relative abundance within the complete 16S and 18S dataset was > 0.01%, respectively. As a result of this the number of ASVs decreased from 7736 to 430 and from 1577 to 58 within the 16S and 18S datasets, respectively. After these filtration steps the final library size comprised 5908-18328 and 2924-417990 reads in 16S and 18S datasets, respectively. Counts of unique ASVs per sample were used to measure richness and Shannońs diversity index was used to combine both richness and evenness – i.e., alpha-diversity [31]. The ‘multcomp’ R package was used to perform *post hoc* multiple comparison test with Tukey’s HSD [32] on alpha-diversity of different sample types between and within T1-crash and T2-healthy using generalized linear models with Poisson distribution. Potential core prokaryotic taxa associated with the focal strain from inoculum to the end of time series were identified applying an occurrence filter with 75% presence threshold separately on T1-crash and T2-healthy 16S dataset.

The compositional differences in prokaryotic and microeukaryotic communities – i.e., beta diversity was measured with unweighted Jaccard distance metric to emphasize presence/absence of community members. Three sample days, both from the beginning and the end of 300 m^3^ bioreactor time series, were included to infer how the bioreactor microbiome composition in the start and end differs and how they relate to the communities in inoculum, intake water and SW bioreactor (pre-filled 300 m^3^ reactor with filtrated seawater). The community differences were visualized in Principal Coordinate Analysis (PCoA) and the pairwise permutational multivariate analysis of variance (PERMANOVA) was performed to test the (dis)similarity associated to each sample type and chosen time series fraction within and between T1-crash and T2-healthy. Beta diversity was also measured via Jaccard distances between T1-crash and T2-healthy bioreactor time series samples, respectively. The Jaccard distances were partitioned to account for the contributions of “nestedness” (species loss) and “turnover” (species gain) towards the measure of total dissimilarity – i.e., *β_Jac_ = β_Jne_ + β_Jtu_* [33, 34]. This was obtained using the R package ‘betapart’ [35]. The associated changes within total beta diversity, turnover, and nestedness, respectively, by sampling day were tested using “adonis” function (PERMANOVA) with 999 permutations from the R package ‘vegań [36]. Taxonomic recruitment of prokaryotes across time series experiments was estimated by calculating the fraction of ASVs that were unobserved at any previous cultivation days [34]. This metric was adapted from the data analysis of a previous study investigating community succession and is available at https://github.com/pnnl/brislawn-2018-founders-species.

### 2.9 Microbiome – Clustering around temporal patterns

The K-medoids clustering method was depolyed following the previously described approach applied to microbiome time series data [37, 38]. It was used to investigate shifts in microbiome structure by examining community members that share similar succession patterns across the crash and healthy time series. Prior to clustering the ASV counts were preprocessed through normalization steps including variance stabilizing transformation (R package ‘DESeq2’) [39], detrending (R package ‘pracmá) [40], and scaling via z-transformation to convert values on z-score scale to ensure intercomparability between ASVs. As the K-medoids is a distance-based clustering method pairwise Euclidean distances were calculated before each ASV was partitioned into K clusters according to similarity criterion using the R package ‘cluster [41]. The clustering efficacy – a balance between number of clusters and data overfitting – was assessed via the Calinski-Harabasz index (Supplementary Figure S1) [42, 43].

### 2.10 Growth conditions of 300 m^3^ bioreactor – Statistics

The Mann-Whitney U test (also known as Wilcoxon rank sum test) was performed to reveal differences in bioreactor physical and biogeochemical conditions between T1-crash and T2-healthy. The potential association between the measured bioreactor conditions and their relationship with identified temporal patterns (cluster medoids) was obtained using Spearman’s rank correlation analysis. Non-parametric analyses were conducted as half of the included factors indicated non-normal distribution based on Shapiro-Wilk normality test (Supplementary Table S1).

### 2.11 Data repository and reproducible analyses

The unprocessed genetic sequencing data for both SSU rRNA 16S and 18S gene amplicons is available in ENA repository under project PRJEB66331. All data including ASV, taxonomy and relative abundance tables, and bioreactor measurements used for analysis and graphing along with R markdown script are available on the Open Science Framework (osf.io) as part of this project: https://osf.io/k2egy/.

## 3 Results

### 3.1 The bioreactoŕs microbiome

Open mass cultivation of marine diatoms within the industrial CCU setting resulted in a complex microbiome that accompanied growth of the target strain *P. glacialis*. The prokaryotic and microeukaryotic community members were diverse and relatively dynamic during both crashed and healthy cultivation periods. This was observed across two time series experiments with lengths of 39 (crashed) and 60 days (healthy), respectively. The first time series – T1-crash – ended with the loss of dominating abundance of the focal strain, *P. glacialis*, as determined by decrease in growth and microscopy observations revealed severe increases of dead and lysed cells. During the second time series from a mass cultivation run – T2-healthy – *P. glacialis* kept its growth and abundance until the operation was ended and biomass was harvested after 60 days. Despite the major differences in conditions and outcomes, the observed richness and diversity, especially for the prokaryotic component of the microbiome, across both time series experiments shared major similarities (Figure 1 and Supplementary Figure S2). The total number of prokaryotic ASVs in T1-crash was 339 and in T2-healthy 368, and the corresponding microeukaryotic richness was 47 and 56 ASVs, respectively. Prokaryotic richness and diversity (Shannon) measured in the 300 m^3^ bioreactor did not differ between the T1-crash and T2-healthy (Tukey’s HSD results given in Supplementary Table S2 and S3), as in total 291 and 256 unique ASVs were observed across time (Figure 1a,b), and the Shannon diversity index ranged from 1.7 to 3.2 in T1-crash and 1.6 to 3.2 in T2-healthy (Supplementary Figure S2). However, only 54-128 ASVs in T1-crash and 49-113 in T2-healthy were present in each sample day, which corresponds to the richness measured within each inoculum, but not within the SW bioreactor (pre-filled 300 m^3^ reactor with filtrated seawater) or intake water samples, which comprised a higher number of ASVs (Figure 1a, b). A clear decrease in Shannon diversity was observed in the middle of T1-crash and T2-healthy. As a somewhat similar pattern was observed in richness, especially in T2-healthy, the change was only partially driven by evenness. The microeukaryotic portion of the bioreactor microbiome (measured via 18S rRNA gene amplicons) showed mainly a contrasting result as compared to the prokaryotic community (Figure 1c, d). The SW reactor and intake water samples comprised a smaller or corresponding number of unique ASVs than the inoculum and per day 300 m^3^ bioreactor samples. Also, the mean number of observed ASVs but not Shannon diversity in the bioreactor across time was significantly different between T1-crash and T2-healthy (Tukey’s HSD, p < 0.001; Supplementary Table S4 and S5). In both time series, the richness and changes that occurred over time were somewhat similar until day 32. Thereafter, the number of observed ASVs stayed constant in T2-healthy, whereas in T1-crash a strong decrease was observed towards the end (Figure 1c, d). This pattern was not inferred from the Shannon diversity index, which instead revealed similar diversity, especially in the end of both time series experiments (Supplementary Figure S2).

**Figure 1.**
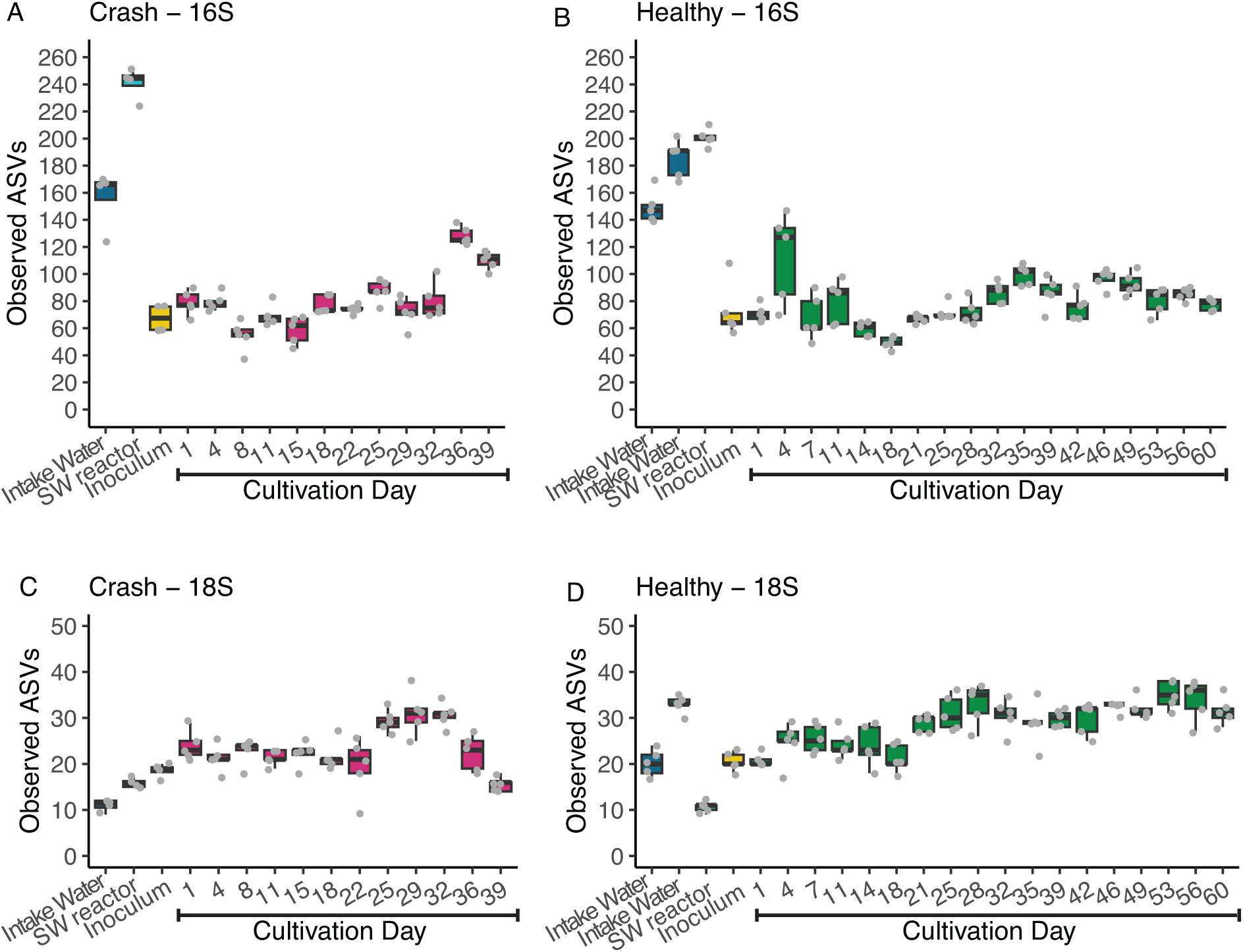
Alpha diversity measured as observed number of unique ASVs in each sample type and across bioreactor time series cultivations. Prokaryotic richness (16S) in time series experiment (A) crash and (B) healthy. Microeukaryotic richnes (18S) in time series experiment (C) crash and (D) healthy. The values of biological replicates (n = 4 or n = 5) are shown over boxplots. Note the different y-axis range between 16S and 18S datasets.

### 3.2 Mass cultivation has an abundant core microbiome

Nearly the same number of ASVs and taxonomic assignments within prokaryotes were found when ASVs were filtered on occurrence (i.e., presence) at a threshold of 75% among the inoculum and 300 m^3^ bioreactor samples. These were 27 and 21 in T1-crash and 25 and 22 in T2-healthy, respectively, that were present throughout the mass cultivation runs. The core ASVs accounted for 20-50% of the unique ASVs observed each sample day in bioreactor of both time series runs. Fourteen of the core ASVs, categorized into nine different bacterial orders, were discovered in the bioreactor and inoculum of the two time series periods. Between 44% and 91% of the prokaryotic community’s daily relative abundance in T1-crash and, correspondingly, between 33% and 91% in T2-healthy, were the taxa that the core ASVs were classified into (core and other ASV(s), henceforth referred to as core taxa) (Figure 2a, b). Supplementary Table S6 contains a list of all the core ASVs together with the corresponding taxonomic assignments.

**Figure 2.**
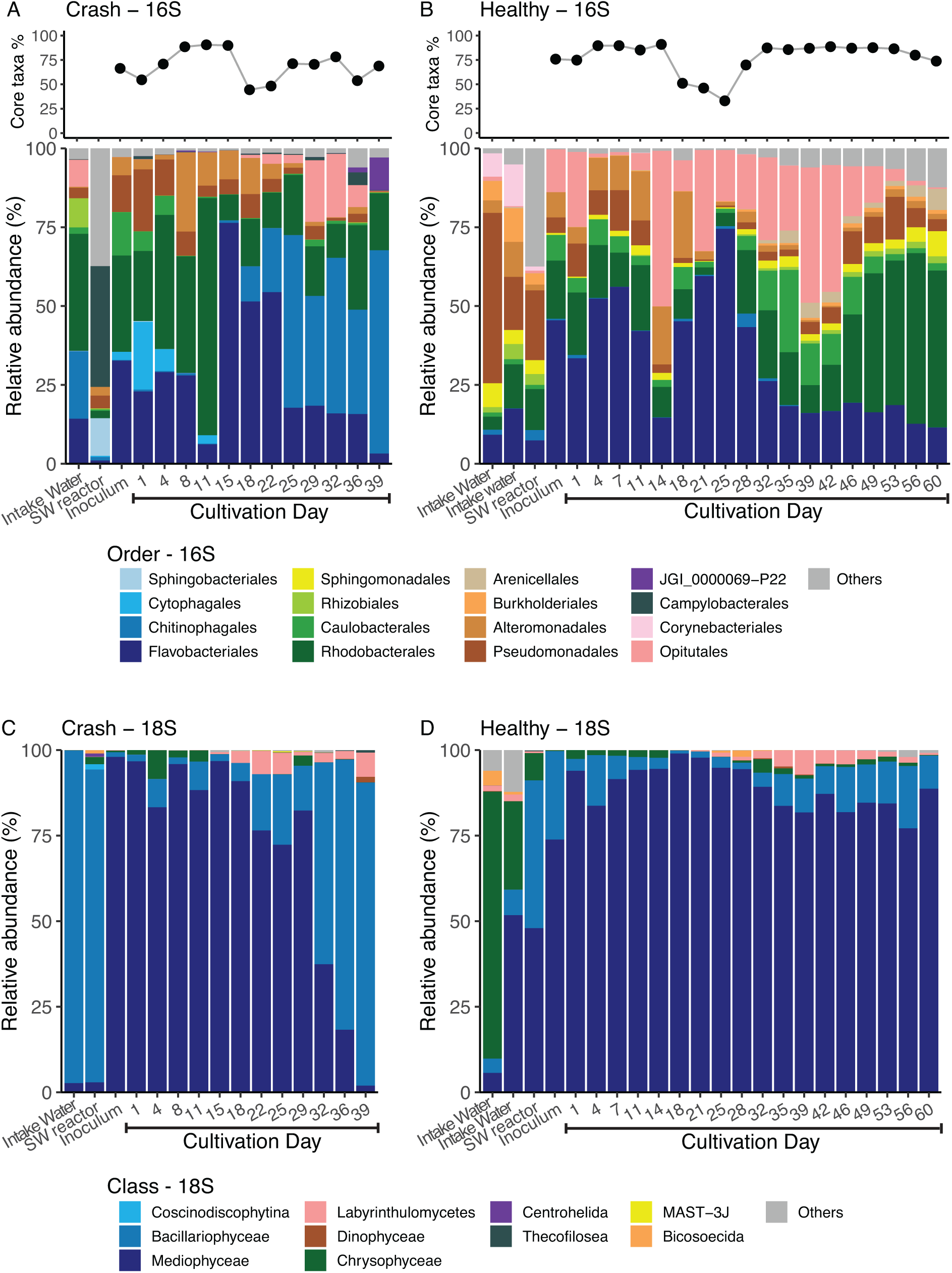
Comparison of community structure in each sample type and across bioreactor time series cultivations. The change in relative abundance of core taxa (core ASVs and other ASVs classified within the same taxonomic assignment) from inoculum to cultivation end and the taxonomic composition of the most abundant prokaryotes classified within taxonomic level order in (A) crash and (B) healthy time series. Taxonomic composition of the most abundant microeukaryotes classified within taxonomic level class in (C) crash and (D) healthy time series.

### 3.3 The reactor’s prokaryotic community structure was distinct between crash versus healthy production

When the 16S ASVs were allocated to the taxonomic level order, the prokaryotic community in both time series studies exhibited a dominance of three groups (Figure 2a, b). Specifically, in the T1-crash, the community was dominated by taxa classified within the order *Rhodobacterales* (22%-75% of the relative abundance) until sample day-11, comprising mainly ASVs assigned to genus *Planktotalea* (Figure 2a). In contrast, T2-healthy, *Rhodobacterales* became the most abundant class in the end of the time series, after sample day 46 (28%-54% of relative abundance) and was dominated by ASVs not assigned to a lower taxonomic level (Figure 2b). Taxa indicative to the order *Flavobacteriales* were predominant, notably ASVs assigned to genera NS9 marine group and *Winogradskyella*, for the first half of the T2-healthy time series with exception of sample day-14, while in T1-crash the predominance of the same genera was shorter and limited in the middle of the time series (Figure 2a, b). The third predominant prokaryotic order was different between the time series experiments as taxa classified within order *Chitinophagales*, mainly ASVs assigned to the family *Saprospiraceae*, became the most abundant in the end of T1-crash (25%-64% of relative abundance) (Figure 2a). In contrast to that in T2-healthy taxa classified as *Opitutales*, comprised almost solely by genus indicative to *Lentimonas*, had a high relative abundance in most of the sample days and was the most abundant on sample day-14 (49% of the relative abundance) and in the middle of the second half of the time series (Figure 2b).

### 3.4 The bioreactor’s microeukaryotic community structure was similar until the culture collapse in T1-crash

The focal strain, *P. glacialis*, is currently classified within class *Mediophyceae* [44], although this classification was not given by the SILVA v138 database as 19 different ASVs indicative to class *Mediophyceae* were assigned under the same annotation (*Mediophyceae*) down to genus level. Among these ASVs, ASV *032943128192fa7f0a17b2d2991a72f4*, comprised 2-3 magnitude higher number of reads compared to the other ASVs. It is therefore considered that most of the relative abundance within class *Mediophyceae* is composed by the focal strain, *P. glacialis* in the 18S microeukaryotic communities.

The community structure of microeukaryotes at the taxonomic level class was similar between T1-crash and T2-healthy samples until a rapid turn from *P. glacialis* dominance to predominance of ASVs classified within the class *Bacillariophyceae* after sample day 29 (59%-89% of relative abundance) in T1-crash (Figure 2c, d). It is notable that members of *Bacillariophyceae* were present from the start in both time series with similar abundances until strong occupation took place in T1-crash whereas in T2-healthy the relative abundance stayed constant until the end of experiment. This contamination was caused by small pennate diatoms, as confirmed by preliminary microscopic detection (Supplementary Figure S3). The most abundant ASV within *Bacillariophyceae* was *f030aa085cd7c8b4bb6538425a95161c*. ASVs assigned to the class *Bacillariophyceae* were also present in the community in the SW bioreactor and intake water samples in the T1-crash, but to a lower degree in the T2-healthy run, where taxa representative of the class *Chrysophyceae* were prevalent, particularly in the intake water. However, *Chrysophyceae* did not show increasing occupation across time in the T2-healthy microbiome (Figure 2c, d).

### 3.5 Beta diversity – microbiome community composition between sample types and across time

The community composition of prokaryotes and microeukaryotes changed from inoculation to the end-point in both time series experiments as inferred from pairwise PERMANOVA comparison of the three first (start) and last (end) sample days (T1-crash: 16S, R^2^ = 0.50 and *p* < 0.01 & 18S, R^2^ = 0.61 and *p* < 0.01; T2-healthy: 16S, R^2^ = 0.61, *p* < 0.01 & 18S, R^2^ = 0.63, *p* < 0.01). The communities within start- and end-time fractions were significantly dissimilar to those obtained in corresponding inoculum, intake water, and SW bioreactor samples and these sample types were also dissimilar in relation to each other (Supplementary Table S7 and S8). The Jaccard dissimilarity distances within 16S and 18S datasets, respectively, were visualized by ordination with PCoA which showed that 39.7% and 44.0% of the total variation among all included prokaryotic and microeukaryotic samples, respectively, were explained by the two first principal coordinate axis (Figure 3). Closer inspection of the ordination results indicated that in both time series experiments, the community composition in the early time series were more similar with the parallel inoculum community, except for microeukaryotes in T1-crash, than in the end as the communities exhibited strong differentiation across time (Figure 3). The communities within SW bioreactor and intake water were clearly distinct from those in inoculum and 300 m^3^ bioreactor samples but also from each other, although the 300 m^3^ bioreactor was pre-filled with intake water (Figure 3).

**Figure 3.**
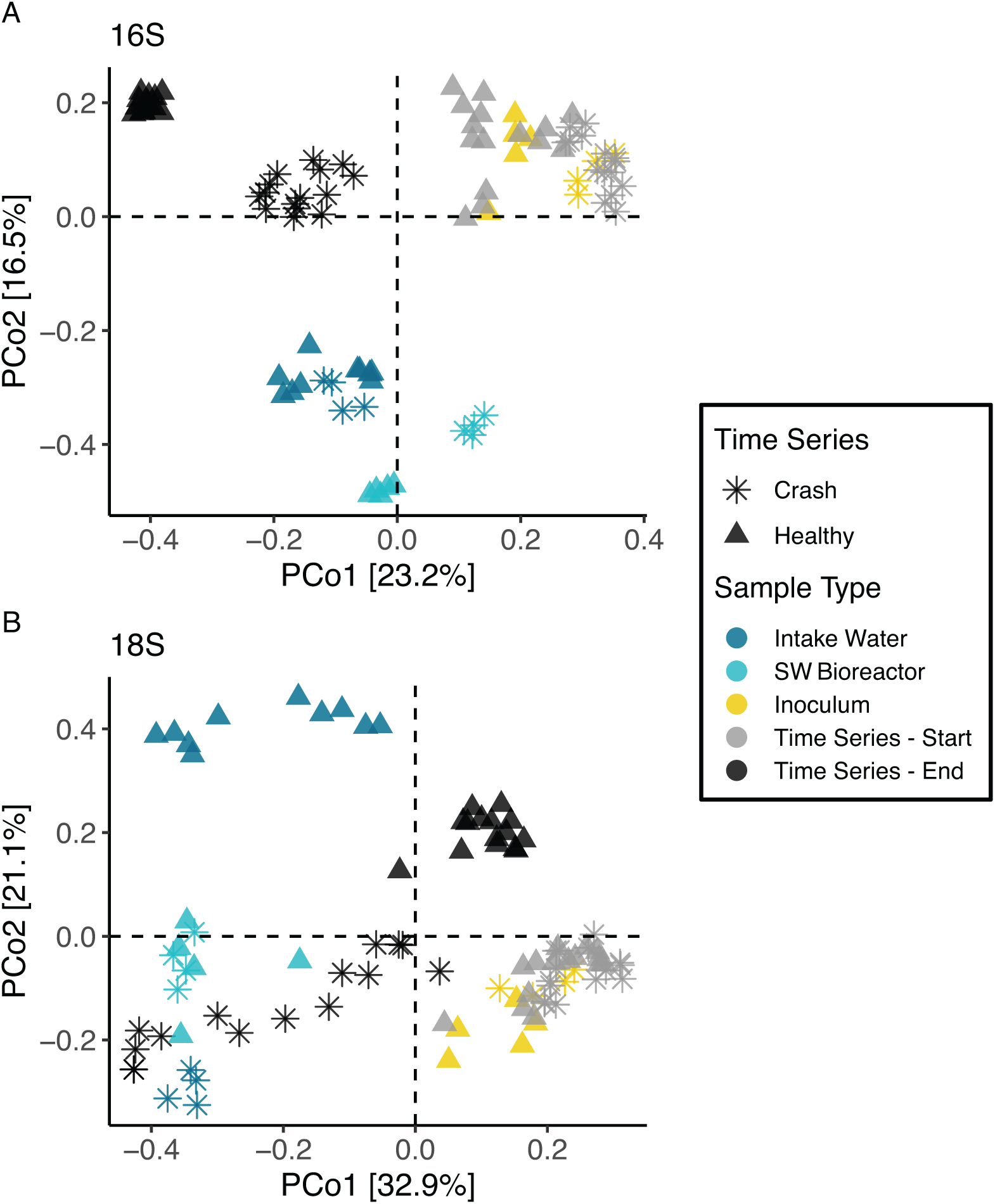
Beta diversity in prokaryotic and microeukaryotic commmunity composition between sample types and bioreactor time series start- and end-time fractions measured with unweighted Jaccard distances and visualized via Principal coordinate analysis. Dissimilarity within (A) prokaryotic and (B) microeukaryotic communities. Time series start-time fraction comprises cultivation days 1, 4 and 8 in crash and 1, 4 and 7 in healthy experiment. The corresponding end-time fractions are 32, 36 and 39 for crash and 53, 56 and 60 for healthy time series.

### 3.6 Beta diversity – contribution of species-loss and replacement to the bioreactoŕs community succession patterns

Temporal variation in the community composition of the bacterial fraction of the microbiome was driven to a greater degree by turnover (species replacement) than nestedness (species loss) in both time series (Figure 4a, b). This inference was obtained by analysis that disentangles the additive contributions of these two antithetic processes to the total beta diversity (as measured by the Jaccard distance) between all-time points across T1-crash and T2-healthy, respectively [33]. The variance attributed to the sampling day was much lower from nestedness than from turnover and total Jaccard in both time series as these explained over 84% and 78% of the observed temporal variation associated with the time points, respectively, in both time series runs (Figure 4 and Supplementary Table S9). Strong turnover is also indicated by the observed prokaryotic richness which stayed rather constant across T1-crash and T2-healthy as under nestedness some level of decrease would be expected through temproal species loss (Figure 1). The bioreactor is exposed to several sources of new taxa that were not continuously monitored in this study, including the input sea water and factory particulate matter from the air. Therefore, the recruitment of prokaryotes was inferred from a metric that indicates the fraction of new taxa introduced to the system at each time point in the time series experiments (Figure 4c). The results revealed that the system is exposed to the detectable seed bank of available community members very quickly. The introduction of previously unobserved ASVs clearly decreased after the 11–14 days of cultivation, and less than 8% and 5% of the ASVs were new in the rest of the cultivation days in T1-crash and T2-healthy, respectively (Figure 4c). Although both of the time series experiments indicated similar pattern, the T1-crash recruited new taxa faster than T2-healthy towards the end of time series.

**Figure 4.**
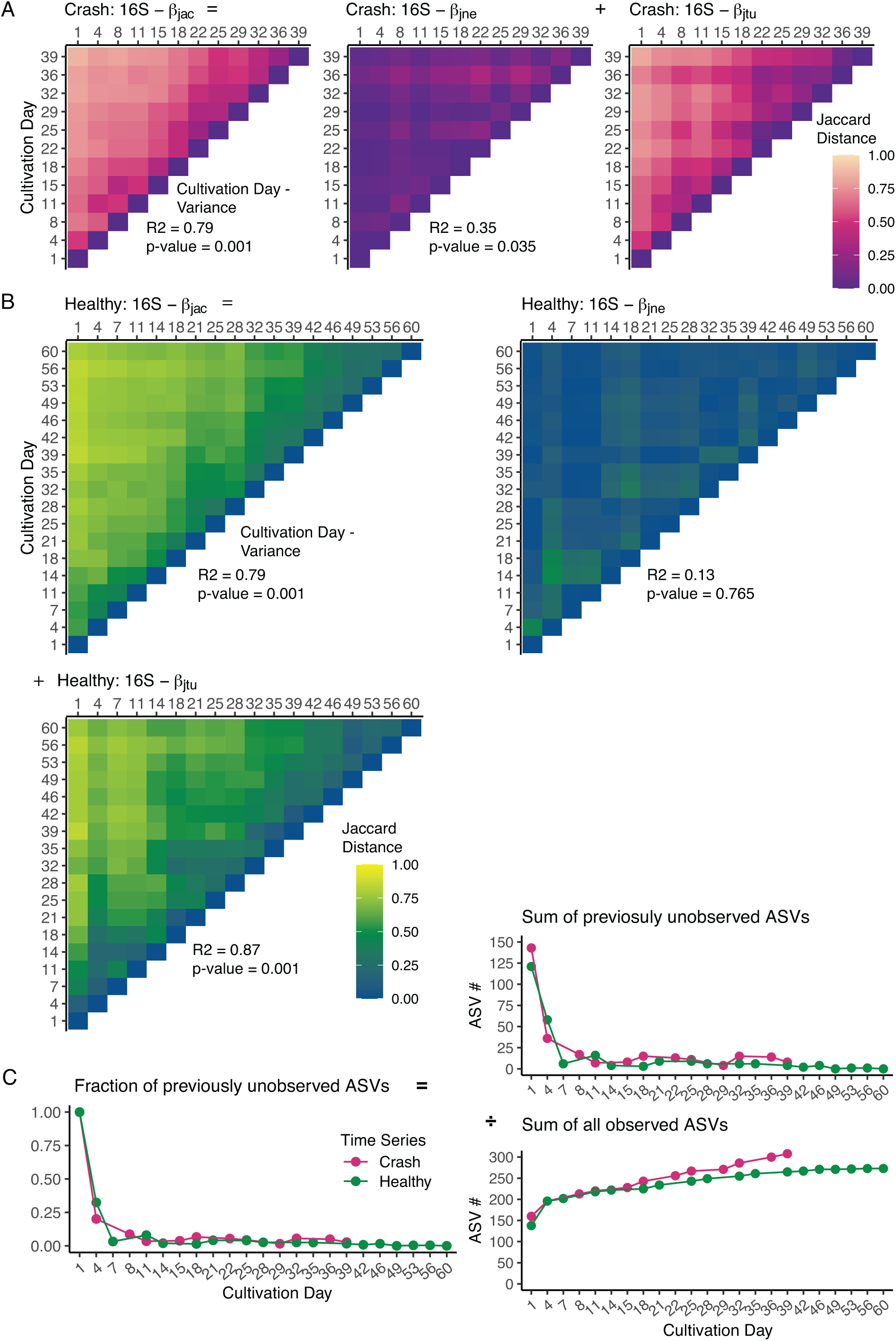
Changes in prokaryotic communities across time. Unweighted Jaccard distance measured between cultivation days in (A) crash and (B) healthy time series experiments. The total Jaccard distance (β_jac_) is partiotioned by contributions from nestedness (β_jne_; species loss) and turnover (β_jtu_; species replacement) i.e., β_jac_ = β_jne_ + β_jtu_. Lighter color represents cultivation days with high dissimilarity. Variance and correspondig significance associated with cultivation day are shown for each process. Introduction of new prokaryotic ASVs over time in crash and healthy time series experiments (C). The fraction of previously unobserved ASVs equals the number of ASVs that were not observed in any of the previous cultivation days and the sum of the total number of ASVs observed at the corresponding time point.

### 3.7 Temporal succession patterns and cooccurring taxa in distinct bioreactor growth environments

The physical and chemical environment in the large 300 m^3^ photobioreactor, as well as cell density of *P. glacialis*, were different between mass cultivation runs. This was established using the Mann-Whitney U test that showed statistically significant differences between *P. glacialis* cell number and raw fluorescence, concentration of inorganic nutrients, DOC, and bioreactor temperature – i.e., most of the monitored factors except pH level (Supplementary Table S10). The measured values of these factors were higher in T2-healthy compared to the T1-crash apart from DOC concentration and bioreactor temperature which were lower (Figure 5 and Supplementary Figure S4).

**Figure 5.**
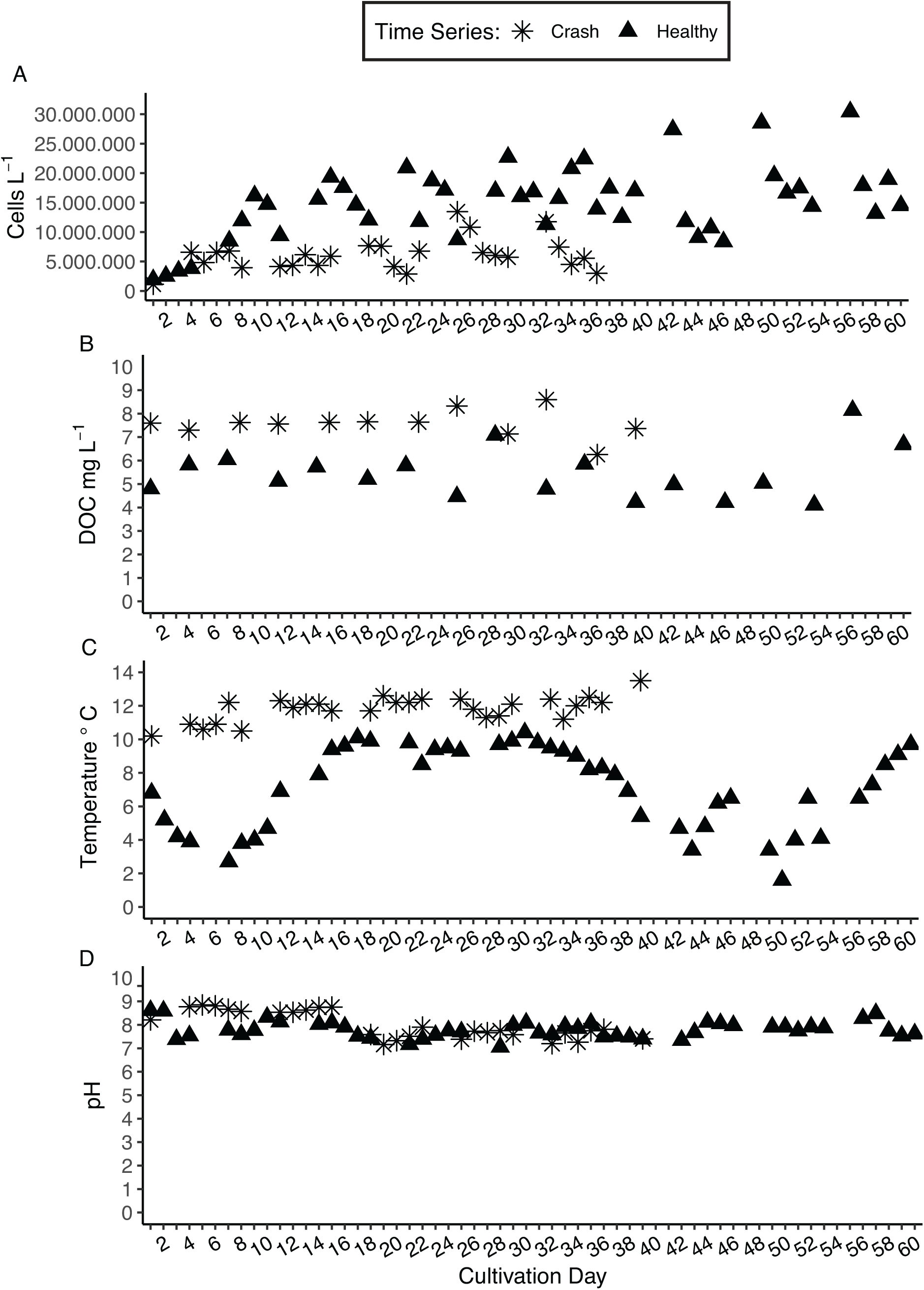
Variation in bioreactor environment across time in crash and healthy cultivation. Daily or twice a week measurements of (A) *P. glacialis* cell density, (B) DOC concentration, (C) bioreactor temperature and (D) pH level. Note that the time series crash was ended after cultivation day-39.

The potential association of community members to the variable bioreactor environment that prevailed across the time series experiments was evidenced through correlation with temporal patterns. A K-medoid clustering algorithm was used to reduce the temporal complexity within the bioreactor microbiome by identifying the major temporal patterns and revealing the related community members within each time series (T1-crash and T2-healthy) and dataset (16S and 18S). The ASVs within both time series and datasets were divided into three clusters based on assessment of the Calinski-Harabasz index which was used as a quality metric (Supplementary Figure S1) [43]. The succession patterns among prokaryotes, as represented by a cluster specific medoid taxon, revealed that part of the community members from both time series datasets (cluster 2 in T1-crash and T2-healthy) exhibited similar temporal dynamics with increase in abundance in the middle of the time series (Figure 6a, b). The medoid of these clusters indicated a significant positive relationship to one of the two factors measuring *P. glacialis* cell density. Yet, the association to the other growth condition factors was not consistent (Figure 6c, d). Also, most of the prokaryotic taxa within cluster 2 (T1-crash and T2-healthy) belonged to the same highly abundant orders *Flavobacteriales*, *Rhodobacterales*, *Opitutales* and *Alteromonadales* (Figure 6e, f). The K-medoid clustering also identified a high number of prokaryotes in T1-crash (cluster 1 and 3) that first decreased and thereafter increased in their abundance towards the culture crash of *P. glacialis* (Figure 6a). Especially in cluster 1 the temporal pattern of medoid showed a strong inverse relationship with *P. glacialis* cell density and DOC concentration and indicated an intensive proliferation of ASVs classified within *Rhodobacterales* and *Chitinophagales* in the last two sample days as these orders comprised the major taxonomic groups within this cluster (Figure 6c, d).

**Figure 6.**
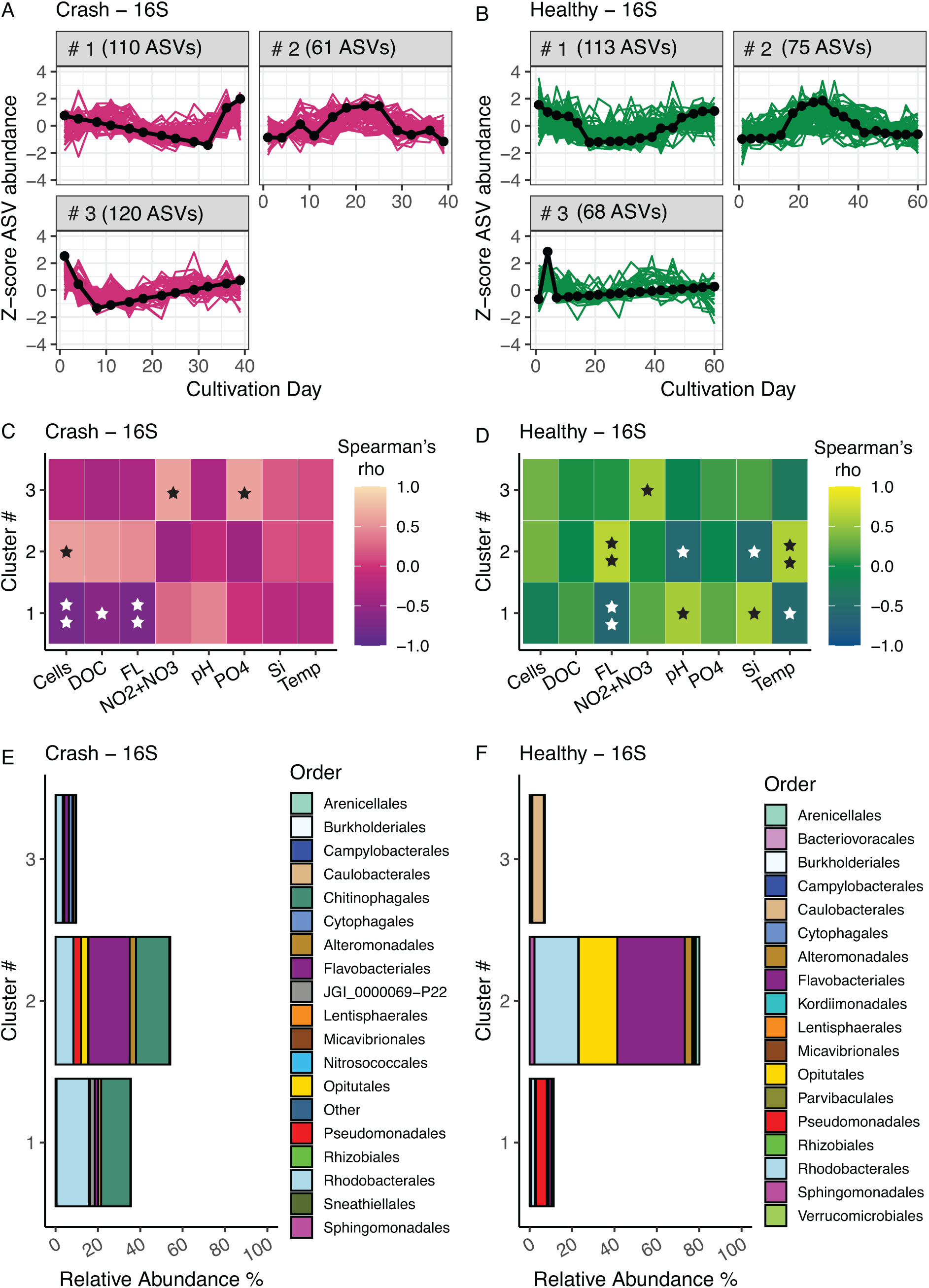
The major temporal succession patterns within prokaryotic communities as assessed via K-medoid clustering analysis with z-transformation and detrending of 16S ASVs. Temproal dynamics of prokaryotic taxa and identified number of ASVs in bioreactor (A) crash and (B) healthy time series experiemnts. The black line indicates cluster specific medoid taxon, a representative for cluster’s temporal pattern. ASV specific z-score is a y-axis and a value 0 denotes the mean abundance. Heatmap of Spearman’s correaltion between temporal dynamics of cluster specific medoid taxon (z-scores) and measured bioreactor factors in time series (C) crash and (D) healthy. The color indicates Spearmna’s rank correlation coefficient (rho) and the significance is denoted with ★(★, p ≤ 0.05; ★★, p ≤ 0.01; ★ ★ ★, p ≤ 0.001). Cells, *P. glacialis* cell density L^-1^; DOC, DOC mg L^-1^; FL, raw fluorescence as a proxy for increase/decrease of *P. glacialis* cell density; NO2-NO3, NO_2_^-^ + NO_3_^-^ µM; pH, pH level; PO4, PO_4_^3-^ µM; Si, Si(OH)_4_ µM, Temp, bioreactor temperature °C. The most common prokaryotic taxa at the taxonomic level order within each cluster and their mean relative abundance in a whole community in time series (E) crash and (F) healthy.

The cluster medoids identified within 18S microeukaryotic community revealed a strong fluctuation in relative abundances and weak correlation to bioreactor conditions across time in both time series (Figure 7a-d). ASVs classified within class *Mediophyceae* and *Bacillariophyceae* were the main groups driving distinct temporal patterns in T1-crash and T2-healthy. In both time series experiments the class *Bacillariophyceae* belonged to a cluster in which the medoid exhibited strong decrease in relative abundance in the middle of time series (Figure 7e, f).

**Figure 7.**
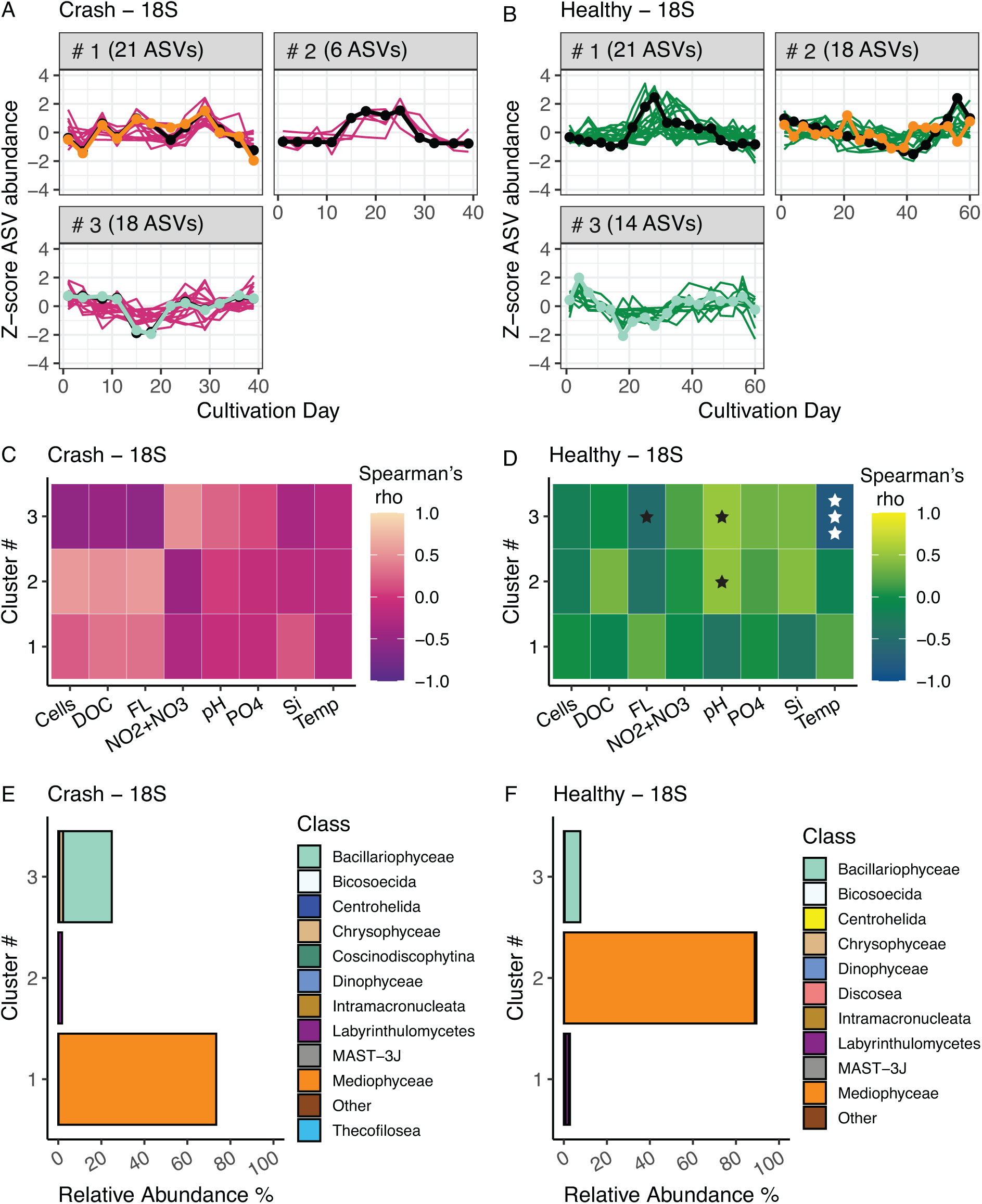
The major temporal succession patterns within microeukaryotic communities as assessed via K-medoid clustering analysis with z-transformation and detrending of 16S ASVs. Temproal dynamics of prokaryotic taxa and identified number of ASVs in bioreactor (A) crash and (B) healthy cultivation experiemnts. The black line indicates cluster specific medoid taxon, a representative for cluster’s temporal pattern. ASV specific z-score is a y-axis and a value 0 denotes the mean abundance. The orange line specifies temporal dynamics of the most abundant ASV classified as *Mediophyceae* representing an expected focal strain, and the light green line specifies the most abundant ASVs classified as *Bacillariophyceae*, representing a potential pennate contaminant. Heatmap of Spearman’s correaltion between temporal dynamics of cluster specific medoid taxon (z-scores) and measured bioreactor factors in time series (C) crash and (D) healthy. The color indicates Spearmna’s rank correlation coefficient (rho) and the significance is denoted with ★(★, p ≤ 0.05; ★★, p ≤ 0.01; ★★★, p ≤ 0.001). Cells, *P. glacialis* cell density L^-1^; DOC, DOC mg L^-1^; FL, raw fluorescence as a proxy for increase/decrease of *P. glacialis* cell density; NO2-NO3, NO_2_^-^ + NO_3_^-^ µM; pH, pH level; PO4, PO_4_^3-^ µM; Si, Si(OH)_4_ µM, Temp, bioreactor temperature °C. The most common microeukaryotic taxa at the taxonomic level class within each cluster and their mean relative abundance in a whole community in time series (E) crash and (F) healthy.

## 4 Discussion

This study was designed to measure and compare the microbial community succession patterns of an industrial scale CCU photobioreactor during both successful and crashed operations. While contamination control is important, it is not usually feasible to operate large scale, open airlift photobioreactors as axenic cultures of microalgae. Hence, industrial scale production of microalgal biomass in open systems is also mass cultivation of the associated microbial community members. This is not necessarily a hindrance to the goals of algal CCU where pure cultures are not required for success, if the bioreactor environment can effectively select for a stable microbial community that promotes growth and abundant proliferation of a focal strain capable of converting CO_2_-enriched flue gas into biomass and bioproducts. This study highlights an example of how a unique core microbiome, associated with cold-water marine environments, stabilizes within a photobioreactor used for mass cultivation of diatoms and how these types of microbiomes represent an important signature and possibly a controlling variable associated with successful operation.

### 4.1 CCU bioreactors are unique industrial marine microbial ecosystems

This study characterized a diverse microbial ecosystem within an industrial scale photobioreactor that was distinct from sea water inflow and inoculum (Figure 2 and 3). This is despite the fact that the operation has been designed to promote growth of large marine diatoms – i.e., *P. glacialis.* The microbial community structure is also sensitive to unique temporal succession patterns that serve as signatures of success or crash. New taxa are recruited to the system via inputs from the semicontinuous inflow of sea water from the adjacent costal bay [45] and likely the air-water exchanges within the industrial environment. The majority of taxonomic recruitment was observed within the prokaryotic portion of the microbiome but observations were also made with respect to microeukaryotic taxa. However, the reactor microbiomés recruitment of new prokaryotic taxa stabilizes very quickly (Figure 4c). Although the bioreactor might be exposed to a larger number of taxa e.g., ASVs removed by the 0.01% abundance threshold or ASVs not detected, these taxa are not taking hold in the community. Changes in community structure were observed on the time scale of days, in a similar fashion to previous observations from temperate and coastal cold-water microbial ecosystems [38, 46]. It is expected that the diverse community comprises members with different ecological properties, which potentially have a strong impact on the function of the production system in relation to CCU [17]. In addition, most of the major bacteria groups *Flavobacteriales*, *Rhodobacterales* and orders within *Gammaproteobacteria* found in the 300 m^3^ bioreactor have frequently been reported to associate with diatom driven phytoplankton blooms in natural ecosystems or in diatom monoculture experiments (Figure 2a, b) [11, 31, 47, 48].

Bacteria that are commonly associated with marine diatoms are known to utilize different alga-derived organic substrates – *i.e.,* DOC that is released across different bloom or growth stages or generated through temporal substrate degradation [46, 49]. These dynamic processes at least partially underpin the successional blooming patterns of the prokaryotic groups that were identified in this study and other investigations [38, 47]. The orders *Opitutales* (*Verrumicrobiota*) and *Chitinophagales* which also belonged to the most abundant community ASVs from the current study are not considered either uncommon or specific to our production system as they have also been documented in environmental and experimental studies. Members within *Opitutales* are likely part of free-living community thriving in eutrophic environments such as wastewater related microalgae cultivations, with a capacity to degrade complex polysaccharides [50]. Members within the *Saprospiraceae* family (*Chitinophagales*) have been reported to interact with diatoms [51].

### 4.2 Importance of a strong core microbiome in mass cultivation

The relatively long and consecutively repeated temporal coverage on microbial community composition provided us an excellent opportunity to infer reoccurring community succession processes. The bioreactor harbors a prominent core microbiome although the taxonomic composition underwent effective replacement of prokaryotes (strong turnover) between time points (Figure 4). We found that the core ASVs comprised a high portion of the total number of unique ASVs observed on each sample day in both time series. Many prokaryotic taxa that are introduced by intake water and the pre-filled bioreactor at the beginning are replaced after a while as they do not integrate into the core community, as evidenced by the strong contribution of turnover towards total community dissimilarity. Consequently, the bioreactor functions as a selective environment because of the very different physical and biogeochemical conditions that these taxa suddenly encounter in comparison to the coastal environment, such as low pH and elevated levels of light, nutrients, and DOC, or because of the core taxa’s competition and antagonistic relationships, which inhibit the growth of the other taxa. It is also expected that *P. glacialis* imposes a selective force that shapes core microbiome as partially evinced by the observed changes in the prokaryotic community when small pennate diatom contaminates supplanted the *P. glacialis* focal strain. The relative strength of diatom-driven selection compared to the physical bioreactor environment remains unclear, although it is understood that the closely associated community is structured by microalgae-specific manner and can persist through environmental variation [12, 52, 53]. The high similarity in the core microbiome between the two cultivation experiments indicates for a certain level of specificity with the production system as it is designed for mass cultivation of *P. glacialis*. It is noted that at least part of the taxa introduced to the bioreactor by intake water influx may reside in the system below detection or were excluded from the analysis along with the applied abundance filter (0.01%). The core taxa (including core and other ASVs classified within the same taxonomic assignment) accounted for an average of 69% and 77% of the relative abundance within bioreactor samples taken from T1-crash and T2-healthy, respectively (Figure 2a, b). Nevertheless, on the majority of sample days, the high abundance was accounted for by a small number of core taxa, whose predominance fluctuated and was only permanent for a short while. Consequently, the temporal change in the prokaryotic community structure was significantly influenced by a small subset of core taxa. The effects of strong temporal variations in community structure and the interactions of the (core) microbiome on the productivity, stability, and/or controllability of the production system are not well understood. Increasing complexity, or the number of interactions within cooccurring relationships, might positively affect stability and, in turn, have an impact on the growth and biomass concentration of the focal strain. [54]. On the other hand, a previous study on the microbiome of marine sponges inferred that a dense core microbiome with few abundant taxa had a stronger positive impact on community level stability than more diverse assemblies of sparsely abundant taxa. [55].

### 4.3 P. glacialis is a promising candidate for mass cultivation

Despite the relatively diversified coinhabiting microbiome, *P. glacialis* grows robustly in the mass cultivation system from inoculation through semicontinuous operation of the 300 m^3^ photobioreactor. While this work is the first to look at the whole community of the bioreactor, mass cultivation of *P. glacialis* for CCU has been successfully demonstrated in industrial scale trials for several years before [5]. Reactor crash events are rare and conversion of industrial CO_2_ to *P. glacialis* biomass is typically successful at scale, regardless of cooccurring microeukaryotes identified in this study. This relatively novel marine diatom is a viable agent for large scale algal CCU and production of high-value biomass [18].

### 4.4 Biological factors associated with reactor crash

Even with a well controlled inoculation and a robust start during mass cultivation, the reactor can occasionally crash. One of the main obstacles when expanding the size of a bioreactor or switching from a closed to an open cultivation system is the high input water volume delivered into the reactor in production systems with (semi)continuous operation, which makes them susceptible to contamination. This is a well-recognized challenge for open cultivation systems even though bioreactor crash events are rarely documented in peer reviewed scientific literature. The research reports on biomass production collapse or culture crash often attribute it to contaminants exhibiting predatory behavior or resource competition [8, 15, 56]. In the present study, the culture collapse in T1-crash was presumably not caused by predation but rather competition or antagonist interactions from another marine diatom or a member from the prokaryotic community. Our study design does not allow for a conclusive mechanistic understanding of the factors that lead to the culture crash; however, our findings suggest two taxa that showed temporal blooming patterns in relation to the observed crash. The *Saprospiraceae* family (order *Chitinophagales*) was the only highly prevalent member of the core community in T1-crash but not in T2-healthy. As algicidal activity has been connected on filamentous members of *Saprospiraceae* [51, 57], the presence of an antagonistic community member against *P. glacalis* should be confirmed in further investigations. We also observed a strongly increasing abundance of pennate diatoms within class Bacillariophyceae towards the end of T1-crash, supported by intake water inflow (Figure 2c), that might have caused competition over resources. Although the cause behind the culture crash remains highly tentative, our findings provide a premise for targeted experiments to uncover taxa with functions that inhibit algal production in this specific industrial system.

### 4.5 Variable environments and blooming patterns within the bioreactor

The large scale mass cultivation system is also unique in relation to the prevailing environment created in the bioreactor and therefore difficult to replicate. This is primarily a result of seasonal variations in the coastal environment and constant production optimization. Although the measured growth conditions varied and were distinct between the time series experiments, the K-medoids clustering did not reveal a clear correlation between the temporal abundance patterns of microeukaryotes and bioreactor factors. Instead, the impact on the prokaryotic community structure might have been stronger (Figure 6). For example, the direct impact of lower and more variable temperature in T2-healthy than in T1-crash on the growth of focal strain was probably minor as *P. glacialis* is shown to obtain a strong growth across the temperature range of 2-12 °C [18]. The growth rate of marine bacteria is known to have a pronounced positive relationship within the similar temperature range [58]. Also, an elevated temperature was reported to increase the abundance of microalgae-associated bacteria and simultaneous carbon assimilation when natural seawater was studied in experimental conditions [59]. Moreover, major differences were observed on the cell density of *P. glacialis* and biogeochemical environment. While the cell density of *P. glacialis* was clearly lower across T1-crash than T2-healthy, the measured DOC concentration revealed the opposite (Figure 5). The dataset does not provide a clear insight on why the DOC level was almost constantly 2.1 mg L^-1^ higher in T1-crash than in T2-healthy or how it potentially governed the cooccurring microbial community. DOC excretion can reflect physiological state of microalgae as a response to stress or it can be triggered by biological interactions [60–62]. The lower cell density in this case does not directly relate to decreased growth (before culture crash) which in turn could indicate stress, because the dilution rate was higher in T1-crash than T2-healthy. However, the concentration of inorganic nutrients, especially silicate in the first half of the time series, was clearly lower in T1-crash than in T2-healthy and therefore might have impacted the cellular response of *P. glacialis* (Supplementary Figure S4).

### 4.6 Core microbiome – an important milestone for further research

This study uncovered the core microbiome associated with mass cultivation of microalgae for CCU and elucidates the role of recurring prokaryotic taxa associated with the open, large scale industrial photobioreactor designed to biomass production of the cold-water diatom *P. glacialis*. The findings of this study show that a stable and recurring core microbiome can significantly influence the productivity and stability of industrial-scale open photobioreactors. The core microbiome, comprising key prokaryotic taxa within orders assigned to *Flavobacteriales*, *Rhodobacterales*, and *Opitutales*, shows resilience and adaptability within the dynamic environment of the bioreactor, suggesting relationships that support the growth of the focal diatom strain, *P. glacialis*. However, the specific functional roles of these core microbial members remain unexplored. Understanding these roles will unlock insights into optimizing bioreactor conditions and enhancing biomass yield and CO_2_ removal from industrial flue gas. Applying targeted experiments using isolation and laboratory coculturing of the members from core community with *P. glacialis* alongside metagenomic and -transcriptomic methods are essential [12, 63]. Also, continuing monitoring the microbiome succession and community profiles over variable production environments and processes enables unraveling the occasional detrimental contaminants in this unique production system. The knowledge will support future research focusing on microbiome engineering to selectively promote beneficial microbial interactions while suppressing antagonistic species that lead to culture crashes [64]. By manipulating the microbial community structure through targeted interventions, it may be possible to achieve higher efficiency and stability in CCU processes, thereby advancing the scalability and economic feasibility of these industrial scale marine diatom cultivation systems.

## 5 Conclusion

This work shows how basic concepts of marine microbial ecology can be used to gain a better understanding of, and perhaps even control over, a promising industrial biotechnology for CCU. Nevertheless, it is difficult to determine the mechanistic influence of the microbiome on large-scale biomass production in large industrial-scale photobioreactors since they are difficult and expensive to modify and operate with high replication for experimental purposes only. Each cultivation run of the large vertical photobioreactor is distinct in terms of the cooccurring microbiome community dynamics and maintained growth conditions throughout the *P. glacialis* production process because of its location within a highly seasonal coastal factory setting. However, the reactor is home to a relatively diverse microbial ecosystem that transforms into a core microbiome via turnover of taxa introduced by the inoculum as well as continuous seawater intake and air exposure to factory particles. Alterations in the core microbiome may influence or result from declining and or stablized growth of *P. glacialis*. Consequently, the unique signature of the reactor’s microbiome is indicative of stable carbon capture and utilization operations, playing a pivotal role in the efficient conversion of industrial CO_2_ into marine biomass.

## Author contributions

HCB and NJA conceptualized the study. IHG, GKE and LI carried out the cultivation and sampling. SK and SP processed molecular samples and amplicon sequence data. NJA and HCB conducted data analysis and wrote and the manuscript draft. All authors contributed to the final version.

## Aknowledgements

This project was made possible through tight collaboration and technical support from Finnfjord AS, with specific acknolwdegements to Hans Christian Eilertsen, Jo Strømholt, John-Steinar Bergum and Geir-Henning Wintervoll.

## Supporting information

upplementary Figures, Tables and Other

