## Supplementary material for "The Microbiome of Bioreactors Containing Mass-Cultivated Marine Diatoms for Industrial Carbon Capture and Utilization": upplementary Figures, Tables and Other

**Affiliations:**
*1. Norwegian College of Fisheries Sciences, UiT - The Arctic University of Norway, Tromsø, Norway*
*2. The Arctic Centre for Sustainable Energy, UiT - The Arctic University of Norway, Tromsø, Norway*

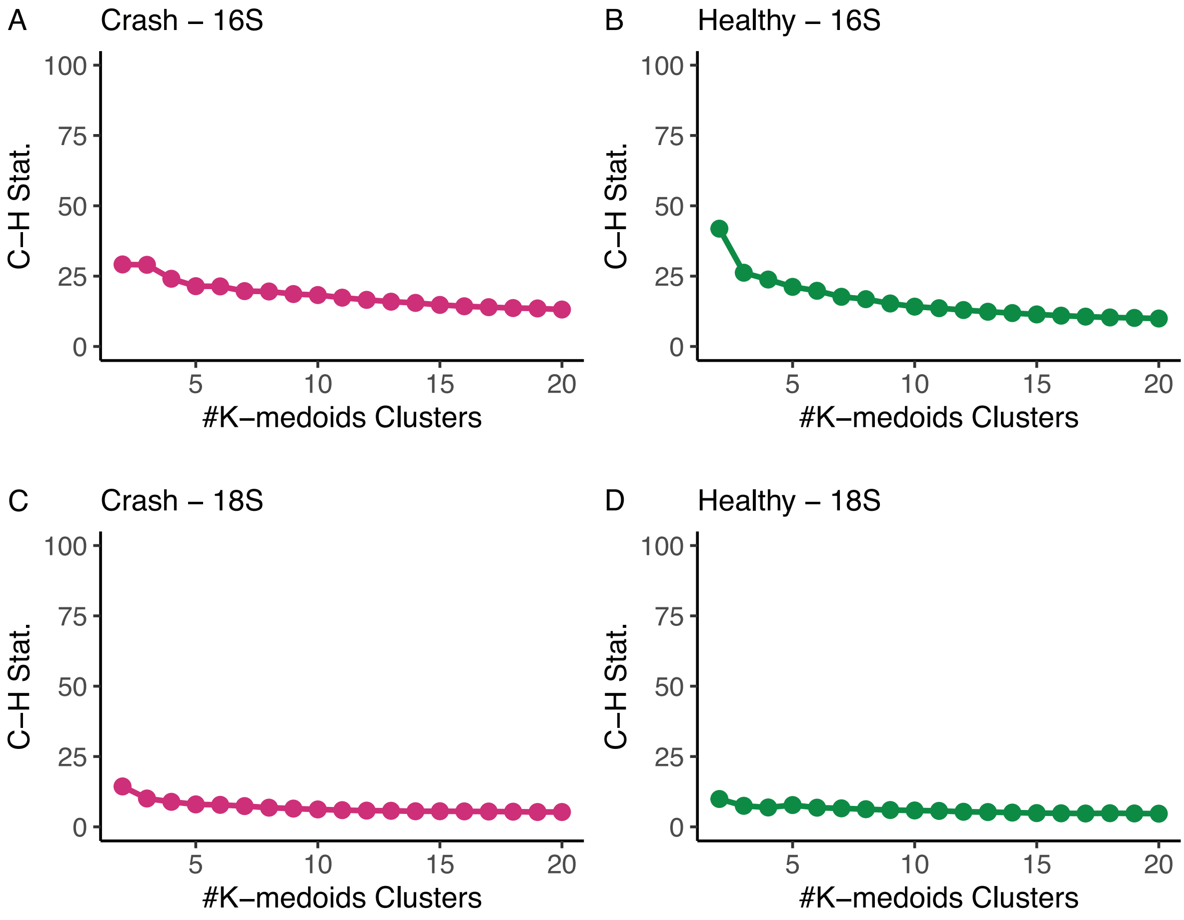


**Supplementary Figure S1.** The Calinski-Harabasz index (y-axis, C-H Stat.) was used to determine the number of K-medoids clusters. C-H Stat is a normalized ratio of the inter and intra cluster variance that was calculated across predefined number of clusters, here 2-20, for each dataset (16S/18S) and time series experiment (crash/healthy). The aim is to obtain a number of clusters whereafter the C-H Stat ratio does not decrease although number of clusters increases. Three clusters were selected to represent the major temporal dynamics of 16S ASVs in (A) crash and (B) healthy time series. The temporal dynamics of 18S ASVs in (C) crash and (D) healthy time series were also characterized with three clusters.

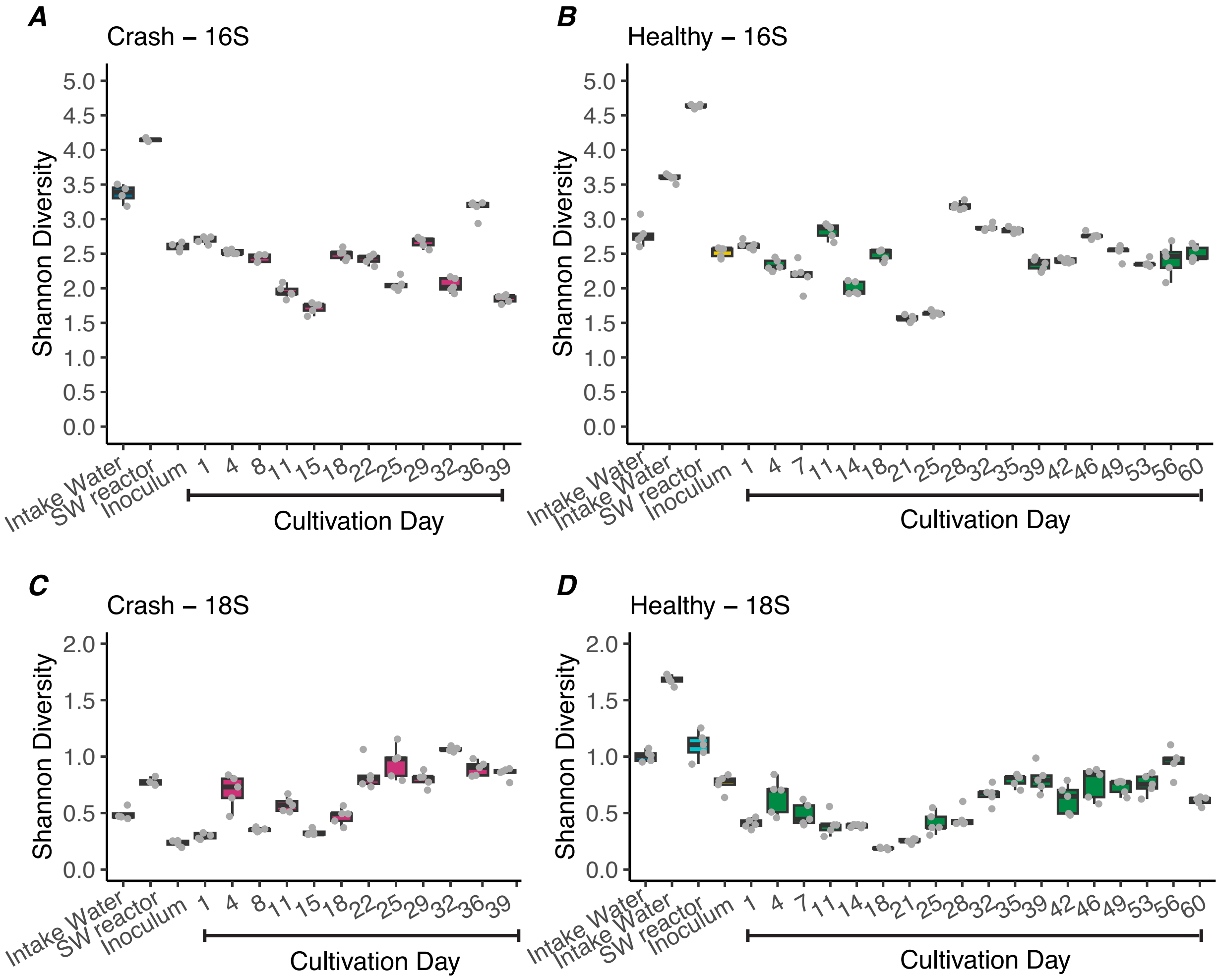


**Supplementary Figure S2.** Alpha diversity measured as Shannon diversity in each sample type and across bioreactor time series cultivations. Prokaryotic diversity (16S) in time series experiment (A) crash and (B) healthy. Microeukaryotic diversity (18S) in time series experiment (C) crash and (D) healthy. The values of biological replicates (n = 4 or n = 5) are shown over boxplots. Note the different y-axis range between 16S and 18S datasets.

**Supplementary Table S1.** Results of Shapiro-Wilk normality test on measured bioreactor factors.

| Factor | statistic | p-value |
|---|---|---|
| DOC mg $L^{-1}$ | 0.931418 | 0.053559 |
| Bioreactor temperature ºC | 0.925641 | 0.000349 |
| pH | 0.943839 | 0.002733 |
| O2 mg $L^{-1}$ | 0.913514 | 0.000965 |
| Raw FL | 0.972742 | 0.114617 |
| Cells $L^{-1}$ | 0.954702 | 0.011207 |
| $NO_2^-$ + $NO_3^-$ µM | 0.915064 | 0.004761 |
| $PO_4^{3-}$ µM | 0.725639 | 2.053780 |
| $Si(OH)_4$ µM | 0.982184 | 0.757734 |

**Supplementary Table S2.** Multiple comparison of prokaryotic (16S dataset) richness (alpha diversity) between sample types and time series – crash vs. healthy – tested with Tukey's HSD using generalized linear models on Poisson distribution. Sample type suffix denotes time series experiment: 1 = Time series crash, 2 = Time series healthy.

```
Simultaneous Tests for General Linear Hypotheses

Multiple Comparisons of Means: Tukey Contrasts


Fit: glm(formula = Value ~ Sample_Type_Time, family = "poisson", data = .)

Linear Hypotheses:
                                       Estimate Std. Error z value Pr(>|z|)
Bioreactor.2 - Bioreactor.1 == 0       -0.01802    0.01860  -0.969  0.97362
Inoculum.1 - Bioreactor.1 == 0         -0.18438    0.06264  -2.944  0.05369 .
Inoculum.2 - Bioreactor.1 == 0         -0.10784    0.05441  -1.982  0.45217
Intake.water.1 - Bioreactor.1 == 0      0.66025    0.04247  15.547  < 0.001 ***
Intake.water.2 - Bioreactor.1 == 0      0.72639    0.02836  25.615  < 0.001 ***
SW.bioreactor.1 - Bioreactor.1 == 0     1.09200    0.03526  30.968  < 0.001 ***
SW.bioreactor.2 - Bioreactor.1 == 0     0.90851    0.03469  26.193  < 0.001 ***
Inoculum.1 - Bioreactor.2 == 0         -0.16636    0.06211  -2.679  0.11016
Inoculum.2 - Bioreactor.2 == 0         -0.08981    0.05380  -1.669  0.67004
Intake.water.1 - Bioreactor.2 == 0      0.67828    0.04168  16.273  < 0.001 ***
Intake.water.2 - Bioreactor.2 == 0      0.74441    0.02717  27.402  < 0.001 ***
SW.bioreactor.1 - Bioreactor.2 == 0     1.11002    0.03431  32.352  < 0.001 ***
SW.bioreactor.2 - Bioreactor.2 == 0     0.92653    0.03372  27.479  < 0.001 ***
Inoculum.2 - Inoculum.1 == 0            0.07655    0.08045   0.951  0.97617
Intake.water.1 - Inoculum.1 == 0        0.84464    0.07290  11.586  < 0.001 ***
Intake.water.2 - Inoculum.1 == 0        0.91077    0.06569  13.864  < 0.001 ***
SW.bioreactor.1 - Inoculum.1 == 0       1.27638    0.06896  18.510  < 0.001 ***
SW.bioreactor.2 - Inoculum.1 == 0       1.09290    0.06866  15.917  < 0.001 ***
Intake.water.1 - Inoculum.2 == 0        0.76809    0.06597  11.643  < 0.001 ***
Intake.water.2 - Inoculum.2 == 0        0.83423    0.05790  14.407  < 0.001 ***
SW.bioreactor.1 - Inoculum.2 == 0       1.19983    0.06158  19.484  < 0.001 ***
SW.bioreactor.2 - Inoculum.2 == 0       1.01635    0.06125  16.593  < 0.001 ***
Intake.water.2 - Intake.water.1 == 0    0.06613    0.04686   1.411  0.82750
SW.bioreactor.1 - Intake.water.1 == 0   0.43174    0.05133   8.411  < 0.001 ***
SW.bioreactor.2 - Intake.water.1 == 0   0.24826    0.05094   4.874  < 0.001 ***
SW.bioreactor.1 - Intake.water.2 == 0   0.36561    0.04044   9.041  < 0.001 ***
SW.bioreactor.2 - Intake.water.2 == 0   0.18212    0.03994   4.560  < 0.001 ***
SW.bioreactor.2 - SW.bioreactor.1 == 0 -0.18348    0.04510  -4.068  0.00104 **
---
Signif. codes:  0 ‘***’ 0.001 ‘**’ 0.01 ‘*’ 0.05 ‘.’ 0.1 ‘ ’ 1
```

**Supplementary Table S3.** Multiple comparison of prokaryotic (16S dataset) Shannon diversity (alpha diversity) between sample types and time series – crash vs. healthy – tested with Tukey's HSD using generalized linear models on Poisson distribution. Sample type suffix denotes time series experiment: 1 = Time series crash, 2 = Time series healthy.

```
         Simultaneous Tests for General Linear Hypotheses

Multiple Comparisons of Means: Tukey Contrasts


Fit: glm(formula = Value ~ Sample_Type_Time, family = "poisson", data = .)

Linear Hypotheses:
                                       Estimate Std. Error z value Pr(>|z|)
Bioreactor.2 - Bioreactor.1 == 0        0.04314    0.10815   0.399   0.9999
Inoculum.1 - Bioreactor.1 == 0          0.10808    0.32130   0.336   1.0000
Inoculum.2 - Bioreactor.1 == 0          0.07714    0.29399   0.262   1.0000
Intake.water.1 - Bioreactor.1 == 0      0.36617    0.28527   1.284   0.8875
Intake.water.2 - Bioreactor.1 == 0      0.31156    0.19621   1.588   0.7249
SW.bioreactor.1 - Bioreactor.1 == 0     0.57461    0.25963   2.213   0.3083
SW.bioreactor.2 - Bioreactor.1 == 0     0.68529    0.22428   3.055   0.0396 *
Inoculum.1 - Bioreactor.2 == 0          0.06494    0.31726   0.205   1.0000
Inoculum.2 - Bioreactor.2 == 0          0.03400    0.28957   0.117   1.0000
Intake.water.1 - Bioreactor.2 == 0      0.32303    0.28071   1.151   0.9340
Intake.water.2 - Bioreactor.2 == 0      0.26842    0.18952   1.416   0.8256
SW.bioreactor.1 - Bioreactor.2 == 0     0.53148    0.25461   2.087   0.3843
SW.bioreactor.2 - Bioreactor.2 == 0     0.64215    0.21845   2.940   0.0555 .
Inoculum.2 - Inoculum.1 == 0           -0.03095    0.41880  -0.074   1.0000
Intake.water.1 - Inoculum.1 == 0        0.25808    0.41272   0.625   0.9981
Intake.water.2 - Inoculum.1 == 0        0.20348    0.35702   0.570   0.9990
SW.bioreactor.1 - Inoculum.1 == 0       0.46653    0.39543   1.180   0.9252
SW.bioreactor.2 - Inoculum.1 == 0       0.57721    0.37318   1.547   0.7508
Intake.water.1 - Inoculum.2 == 0        0.28903    0.39183   0.738   0.9947
Intake.water.2 - Inoculum.2 == 0        0.23443    0.33265   0.705   0.9960
SW.bioreactor.1 - Inoculum.2 == 0       0.49748    0.37358   1.332   0.8668
SW.bioreactor.2 - Inoculum.2 == 0       0.60815    0.34994   1.738   0.6241
Intake.water.2 - Intake.water.1 == 0   -0.05460    0.32496  -0.168   1.0000
SW.bioreactor.1 - Intake.water.1 == 0   0.20845    0.36675   0.568   0.9990
SW.bioreactor.2 - Intake.water.1 == 0   0.31912    0.34264   0.931   0.9789
SW.bioreactor.1 - Intake.water.2 == 0   0.26305    0.30271   0.869   0.9859
SW.bioreactor.2 - Intake.water.2 == 0   0.37373    0.27300   1.369   0.8494
SW.bioreactor.2 - SW.bioreactor.1 == 0  0.11067    0.32161   0.344   1.0000
---
Signif. codes:  0 ‘***’ 0.001 ‘**’ 0.01 ‘*’ 0.05 ‘.’ 0.1 ‘ ’ 1
(Adjusted p values reported -- single-step method)
```

**Supplementary Table S4.** Multiple comparison of microeukaryotic (18S dataset) richness (alpha diversity) between sample types and time series – crash vs. healthy – tested with Tukey's HSD using generalized linear models on Poisson distribution. Sample type suffix denotes time series experiment: 1 = Time series crash, 2 = Time series healthy.

```
Simultaneous Tests for General Linear Hypotheses

Multiple Comparisons of Means: Tukey Contrasts


Fit: glm(formula = Value ~ Sample_Type_Time, family = "poisson", data = .)

Linear Hypotheses:
                                       Estimate Std. Error z value Pr(>|z|)    
Bioreactor.2 - Bioreactor.1 == 0        0.20280    0.03313   6.121  < 0.001 ***
Inoculum.1 - Bioreactor.1 == 0         -0.23710    0.11927  -1.988  0.43433    
Inoculum.2 - Bioreactor.1 == 0         -0.12958    0.10208  -1.269  0.88606    
Intake.water.1 - Bioreactor.1 == 0     -0.75698    0.15309  -4.944  < 0.001 ***
Intake.water.2 - Bioreactor.1 == 0      0.12604    0.06686   1.885  0.50504    
SW.bioreactor.1 - Bioreactor.1 == 0    -0.39803    0.12878  -3.091  0.03314 *  
SW.bioreactor.2 - Bioreactor.1 == 0    -0.79402    0.13992  -5.675  < 0.001 ***
Inoculum.1 - Bioreactor.2 == 0         -0.43990    0.11790  -3.731  0.00388 ** 
Inoculum.2 - Bioreactor.2 == 0         -0.33238    0.10048  -3.308  0.01669 *  
Intake.water.1 - Bioreactor.2 == 0     -0.95978    0.15203  -6.313  < 0.001 ***
Intake.water.2 - Bioreactor.2 == 0     -0.07676    0.06439  -1.192  0.91556    
SW.bioreactor.1 - Bioreactor.2 == 0    -0.60083    0.12751  -4.712  < 0.001 ***
SW.bioreactor.2 - Bioreactor.2 == 0    -0.99682    0.13876  -7.184  < 0.001 ***
Inoculum.2 - Inoculum.1 == 0            0.10752    0.15239   0.706  0.99560    
Intake.water.1 - Inoculum.1 == 0       -0.51988    0.19037  -2.731  0.09149 .  
Intake.water.2 - Inoculum.1 == 0        0.36314    0.13143   2.763  0.08430 .  
SW.bioreactor.1 - Inoculum.1 == 0      -0.16093    0.17142  -0.939  0.97605    
SW.bioreactor.2 - Inoculum.1 == 0      -0.55692    0.17995  -3.095  0.03266 *  
Intake.water.1 - Inoculum.2 == 0       -0.62740    0.18010  -3.484  0.00920 ** 
Intake.water.2 - Inoculum.2 == 0        0.25562    0.11605   2.203  0.30072    
SW.bioreactor.1 - Inoculum.2 == 0      -0.26845    0.15994  -1.678  0.65048    
SW.bioreactor.2 - Inoculum.2 == 0      -0.66444    0.16905  -3.931  0.00160 ** 
Intake.water.2 - Intake.water.1 == 0    0.88302    0.16275   5.426  < 0.001 ***
SW.bioreactor.1 - Intake.water.1 == 0   0.35895    0.19647   1.827  0.54577    
SW.bioreactor.2 - Intake.water.1 == 0  -0.03704    0.20395  -0.182  1.00000    
SW.bioreactor.1 - Intake.water.2 == 0  -0.52407    0.14012  -3.740  0.00358 ** 
SW.bioreactor.2 - Intake.water.2 == 0  -0.92006    0.15042  -6.116  < 0.001 ***
SW.bioreactor.2 - SW.bioreactor.1 == 0 -0.39599    0.18639  -2.125  0.34643    
---
Signif. codes:  0 ‘***’ 0.001 ‘**’ 0.01 ‘*’ 0.05 ‘.’ 0.1 ‘ ’ 1
(Adjusted p values reported -- single-step method)
```

**Supplementary Table S5.** Multiple comparison of microeukaryotic (18S dataset) Shannon diversity (alpha diversity) between sample types and time series – crash vs. healthy – tested with Tukey's HSD using generalized linear models on Poisson distribution. Sample type suffix denotes time series experiment: 1 = Time series crash, 2 = Time series healthy.

```
         Simultaneous Tests for General Linear Hypotheses

Multiple Comparisons of Means: Tukey Contrasts


Fit: glm(formula = Value ~ Sample_Type_Time, family = "poisson", data = .)

Linear Hypotheses:
                                       Estimate Std. Error z value Pr(>|z|)
Bioreactor.2 - Bioreactor.1 == 0       -0.16592    0.20970  -0.791   0.9914
Inoculum.1 - Bioreactor.1 == 0         -1.06386    1.04604  -1.017   0.9635
Inoculum.2 - Bioreactor.1 == 0          0.12023    0.53523   0.225   1.0000
Intake.water.1 - Bioreactor.1 == 0     -0.32386    0.73138  -0.443   0.9998
Intake.water.2 - Bioreactor.1 == 0      0.68380    0.31485   2.172   0.3194
SW.bioreactor.1 - Bioreactor.1 == 0     0.13668    0.58872   0.232   1.0000
SW.bioreactor.2 - Bioreactor.1 == 0     0.48451    0.45444   1.066   0.9530
Inoculum.1 - Bioreactor.2 == 0         -0.89793    1.04353  -0.860   0.9858
Inoculum.2 - Bioreactor.2 == 0          0.28615    0.53031   0.540   0.9992
Intake.water.1 - Bioreactor.2 == 0     -0.15793    0.72779  -0.217   1.0000
Intake.water.2 - Bioreactor.2 == 0      0.84973    0.30641   2.773   0.0817 .
SW.bioreactor.1 - Bioreactor.2 == 0     0.30261    0.58425   0.518   0.9994
SW.bioreactor.2 - Bioreactor.2 == 0     0.65044    0.44864   1.450   0.7988
Inoculum.2 - Inoculum.1 == 0            1.18408    1.15388   1.026   0.9616
Intake.water.1 - Inoculum.1 == 0        0.74000    1.25694   0.589   0.9986
Intake.water.2 - Inoculum.1 == 0        1.74766    1.06963   1.634   0.6832
SW.bioreactor.1 - Inoculum.1 == 0       1.20054    1.17964   1.018   0.9633
SW.bioreactor.2 - Inoculum.1 == 0       1.54837    1.11870   1.384   0.8344
Intake.water.1 - Inoculum.2 == 0       -0.44408    0.87873  -0.505   0.9995
Intake.water.2 - Inoculum.2 == 0        0.56358    0.57998   0.972   0.9716
SW.bioreactor.1 - Inoculum.2 == 0       0.01646    0.76409   0.022   1.0000
SW.bioreactor.2 - Inoculum.2 == 0       0.36429    0.66616   0.547   0.9992
Intake.water.2 - Intake.water.1 == 0    1.00766    0.76474   1.318   0.8667
SW.bioreactor.1 - Intake.water.1 == 0   0.46054    0.91230   0.505   0.9995
SW.bioreactor.2 - Intake.water.1 == 0   0.80837    0.83200   0.972   0.9716
SW.bioreactor.1 - Intake.water.2 == 0  -0.54712    0.62968  -0.869   0.9850
SW.bioreactor.2 - Intake.water.2 == 0  -0.19929    0.50639  -0.394   0.9999
SW.bioreactor.2 - SW.bioreactor.1 == 0  0.34783    0.70985   0.490   0.9996
---
Signif. codes:  0 '***' 0.001 '**' 0.01 '*' 0.05 '.' 0.1 ' ' 1
(Adjusted p values reported -- single-step method)
```

**Supplementary Table S6.** Core ASVs and associated taxonomic assignments were identified using a 75% presence threshold within inoculum and bioreactor time series samples. The bolded ASVs occurred both in T1-crash and T2-healthy time series experiments.

| ASV | Phylum | Class | Order | Family | Genus | Species |
|---|---|---|---|---|---|---|
| | | | **T1-Crash** | | | |
| 5a25d986b213abcd a4794f700db75bba | Bacteroidota | Bacteroidia | Chitinophagales | Saprospiraceae | | |
| 6f4e8d53540daa89 7094612f2e212f68 | Bacteroidota | Bacteroidia | Chitinophagales | Saprospiraceae | | |
| d8169705948a3089 52da9fa86125f7cb | Bacteroidota | Bacteroidia | Flavobacteriales | Cryomorphaceae | Vicingus | |
| 0f46c35d6629feaf fa5e8f4b256e255b | Bacteroidota | Bacteroidia | Flavobacteriales | Flavobacteriaceae | NS3a marine group | |
| 2190dbbf36ecbe4b c58d62a9799c4c80 | Bacteroidota | Bacteroidia | Flavobacteriales | Flavobacteriaceae | Ulvibacter | Ulvibacter antarcticus |
| **78d9781cf13aac4c b17acccb5114ccbb** | Bacteroidota | Bacteroidia | Flavobacteriales | Flavobacteriaceae | Ulvibacter | Ulvibacter antarcticus |
| **250dfc56f1cbeb2fe 284b54de7e533c3** | Bacteroidota | Bacteroidia | Flavobacteriales | NS9 marine group | NS9 marine group | |
| **2e0b607baace5a65 97f0e2b60f7c5092** | Proteobacteria | Alpha-proteobacteria | Caulobacterales | Hyphomonadaceae | Algimonas | marine bacterium |
| **6ebf245c08769315 8dd26bfd5b0db63e** | Proteobacteria | Alpha-proteobacteria | Rhodobacterales | Rhodobacteraceae | Planktotalea | |
| **4eb72ad317b099b5 7c248c458eab626b** | Proteobacteria | Alpha-proteobacteria | Rhodobacterales | Rhodobacteraceae | Roseovarius | |
| 40d6343cb5340095 1e705a6e0e4216ad | Proteobacteria | Alpha-proteobacteria | Rhodobacterales | Rhodobacteraceae | Sulfitobacter | |
| 1a04461b51bc23b4 8c24529fbdb95d2b | Proteobacteria | Alpha-proteobacteria | Rhodobacterales | Rhodobacteraceae | Sulfitobacter | |
| **7f821b26f148d936 26119f434dd5b5a5** | Proteobacteria | Alpha-proteobacteria | Rhodobacterales | Rhodobacteraceae | Sulfitobacter | |
| b6d52741562c126a fe54b50139400044 | Proteobacteria | Alpha-proteobacteria | Rhodobacterales | Rhodobacteraceae | Sulfitobacter | |
| ebf00dd6a9329614 e82c0a214cb16490 | Proteobacteria | Alpha-proteobacteria | Rhodobacterales | Rhodobacteraceae | Sulfitobacter | |
| **c197bfccc6da06a3 21f59bc2c96c7162** | Proteobacteria | Alpha-proteobacteria | Sphingo-monadales | Sphingo-monadaceae | Sphingorhabdus | |
| **cd79edc57bfd3c04 cca181d3fea42156** | Proteobacteria | Gamma-proteobacteria | Burkholderiales | Methylophilaceae | Methylotenera | |
| 6c6ffa8f21150c46 fa47c3aaf914b3fb | Proteobacteria | Gamma-proteobacteria | Alteromonadales | Alteromonadaceae | Glaciecola | |
| f9a1f200e9e88d0a 15469c60357e8291 | Proteobacteria | Gamma-proteobacteria | Alteromonadales | Alteromonadaceae | Paraglaciecola | Paraglaciecola arctica |
| **1d52609177ce303a 019c3ee3f9142239** | Proteobacteria | Gamma-proteobacteria | Alteromonadales | Colwelliaceae | Colwellia | |
| **d7c175c5b206160e 3c14ffd3ae2019d7** | Proteobacteria | Gamma-proteobacteria | Alteromonadales | Pseudo-alteromonadaceae | Pseudo-alteromonas | |
| **6d44145ac74fc30f 6713c8ab71a27c96** | Proteobacteria | Gamma-proteobacteria | Nitrosococcales | Methylophagaceae | Methylophaga | |
| **55f551c495c9fbdf4 e7605b2c22444bb** | Proteobacteria | Gamma-proteobacteria | Pseudomonadales | Marinomonadaceae | Marinomonas | |
| cb57ce02d911c1b3 270f8186d5963213 | Proteobacteria | Gamma-proteobacteria | Pseudomonadales | Porticoccaceae | Porticoccus | |
| 0b59a39511050d6a f174534cb1465f28 | Proteobacteria | Gamma-proteobacteria | Pseudomonadales | Pseudo-hongiellaceae | Pseudo-hongiella | |
| **b0c2d8dfce86fc0d 49ed2f43eb3a4955** | Proteobacteria | Gamma-proteobacteria | Pseudomonadales | Spongiibacteraceae | BD1-7 clade | |
| **fc27c909d92c3000 0f2fce40bdda1cf7** | Verruco-microbiota | Verruco-microbiae | Opitutales | Puniceicoccaceae | Lentimonas | |
| | | | **T2-Healthy** | | | |
| 0678751fed080c75 058c56eccca734af | Bacteroidota | Bacteroidia | Flavobacteriales | Crocinitomicaceae | Crocinitomix | |
| 444a5d2e1edf7782 09fd4837af7d2e65 | Bacteroidota | Bacteroidia | Flavobacteriales | Flavobacteriaceae | | |
| 61771b37cea411a6 c3f65546fcb0d14d | Bacteroidota | Bacteroidia | Flavobacteriales | Flavobacteriaceae | | |
| 09a6529ddf88c18b 010c5ec5bf9f289a | Bacteroidota | Bacteroidia | Flavobacteriales | Flavobacteriaceae | Arenibacter | |

| | | | | | | |
|---|---|---|---|---|---|---|
| 95e486ecf1b6a809 d41a224b53ed8cf5 | Bacteroidota | Bacteroidia | Flavobacteriales | Flavobacteriaceae | Maribacter | Bacteroidetes bacterium |
| **78d9781cf13aac4c b17acccb5114ccbb** | Bacteroidota | Bacteroidia | Flavobacteriales | Flavobacteriaceae | Ulvibacter | Ulvibacter antarcticus |
| **250dfc56f1cbeb2fe 284b54de7e533c3** | Bacteroidota | Bacteroidia | Flavobacteriales | NS9 marine group | NS9 marine group | |
| **2e0b607baace5a65 97f0e2b60f7c5092** | Proteobacteria | Alpha-proteobacteria | Caulobacterales | Hyphomonadaceae | Algimonas | marine bacterium |
| 550411d8a864933d a10a00714d167c9e | Proteobacteria | Alpha-proteobacteria | Parvibaculales | | | |
| 28e67f2c5608db5aa 4ea08dc98db352e | Proteobacteria | Alpha-proteobacteria | Rhodobacterales | Rhodobacteraceae | | |
| 9d4259355f60e886 fedb1b462edce1ad | Proteobacteria | Alpha-proteobacteria | Rhodobacterales | Rhodobacteraceae | | |
| **6ebf245c08769315 8dd26bfd5b0db63e** | Proteobacteria | Alpha-proteobacteria | Rhodobacterales | Rhodobacteraceae | Planktotalea | |
| ce888d07bfd01d90 6d0efcb107363d11 | Proteobacteria | Alpha-proteobacteria | Rhodobacterales | Rhodobacteraceae | Pseudo-phaeobacter | |
| **4eb72ad317b099b5 7c248c458eab626b** | Proteobacteria | Alpha-proteobacteria | Rhodobacterales | Rhodobacteraceae | Roseovarius | |
| **7f821b26f148d936 26119f434dd5b5a5** | Proteobacteria | Alpha-proteobacteria | Rhodobacterales | Rhodobacteraceae | Sulfitobacter | |
| **c197bfccc6da06a3 21f59bc2c96c7162** | Proteobacteria | Alpha-proteobacteria | Sphingo-monadales | Sphingomonadaceae | Sphingorhabdus | |
| **cd79edc57bfd3c04 cca181d3fea42156** | Proteobacteria | Gamma-proteobacteria | Burkholderiales | Methylophilaceae | Methylotenera | |
| **1d52609177ce303a 019c3ee3f9142239** | Proteobacteria | Gamma-proteobacteria | Alteromonadales | Colwelliaceae | Colwellia | |
| e3bd4c321c5364d0 1871ebb503a1dc7e | Proteobacteria | Gamma-proteobacteria | Alteromonadales | Colwelliaceae | Colwellia | |
| **d7c175c5b206160e 3c14ffd3ae2019d7** | Proteobacteria | Gamma-proteobacteria | Alteromonadales | Pseudo-alteromonadaceae | Pseudo-alteromonas | |
| **6d44145ac74fc30f 6713c8ab71a27c96** | Proteobacteria | Gamma-proteobacteria | Nitrosococcales | Methylophagaceae | Methylophaga | |
| **55f551c495c9fbdf4 e7605b2c22444bb** | Proteobacteria | Gamma-proteobacteria | Pseudo-monadales | Marinomonadaceae | Marinomonas | |
| e0c5352d6d5c234a d58c8a373d29ca55 | Proteobacteria | Gamma-proteobacteria | Pseudo-monadales | Porticoccaceae | Porticoccus | |
| **b0c2d8dfce86fc0d 49ed2f43eb3a4955** | Proteobacteria | Gamma-proteobacteria | Pseudo-monadales | Spongiibacteraceae | BD1-7_clade | |
| **fc27c909d92c3000 0f2fce40bdda1cf7** | Verruco-microbiota | Verruco-microbiae | Opitutales | Puniceicoccaceae | Lentimonas | |

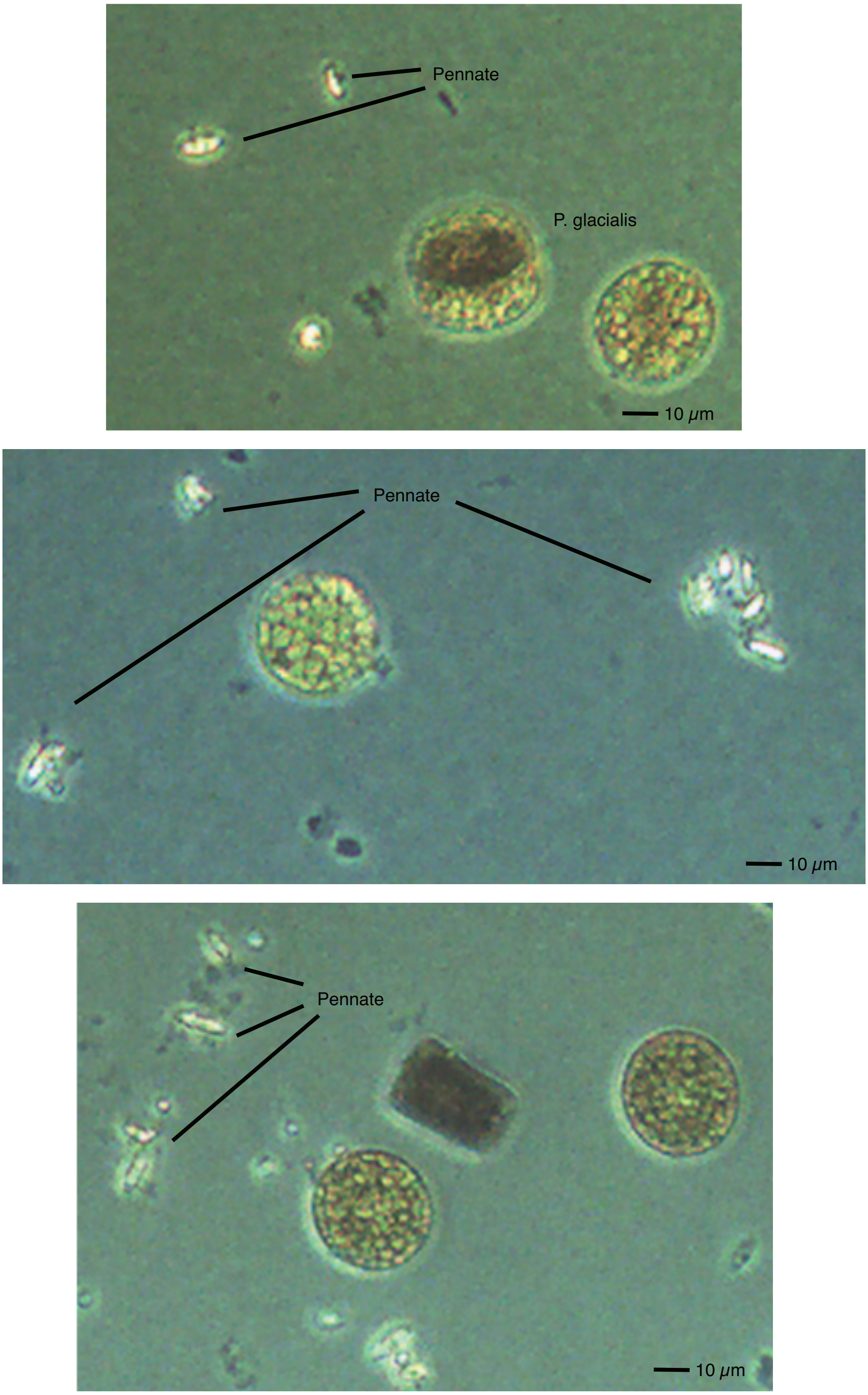


**Supplementary Figure S3.** Observations of pennate contamination in the T1-crash time series experiment under light microscopy examination of culture conditions.

**Supplementary Table S7.** Results of PERMANOVA pairwise comparison generated through 999 permutations between sample types and start- and end-time fraction of prokaryotic (16S) communities in time series healthy and crash. Unweighted Jaccard was used as a distance metric. Sample type suffix denotes time series experiment: 1 = Time series crash, 2 = Time series healthy.

| Groups: 16S dataset | measure | F | R2 | p.value | p.adjusted | Significance |
|---|---|---|---|---|---|---|
| T2.start vs Intake.water.2 | jaccard | 20.1488529 | 0.4669615 | 0.001 | 0.00166667 | ** |
| T2.start vs T2.end | jaccard | 44.1465116 | 0.61190085 | 0.001 | 0.00166667 | ** |
| T2.start vs SW.bioreactor.1 | jaccard | 14.9735065 | 0.4683098 | 0.002 | 0.00264706 | ** |
| T2.start vs Inoculum.1 | jaccard | 9.64485312 | 0.36197809 | 0.001 | 0.00166667 | ** |
| T2.start vs T1.start | jaccard | 13.6241546 | 0.32731367 | 0.001 | 0.00166667 | ** |
| T2.start vs T1.end | jaccard | 29.7962419 | 0.51553943 | 0.001 | 0.00166667 | ** |
| T2.start vs Intake.water.1 | jaccard | 13.9677162 | 0.45104121 | 0.001 | 0.00166667 | ** |
| T2.start vs Inoculum.2 | jaccard | 4.9365174 | 0.21522524 | 0.001 | 0.00166667 | ** |
| T2.start vs SW.bioreactor.2 | jaccard | 20.7459414 | 0.53543521 | 0.002 | 0.00264706 | ** |
| Intake.water.2 vs T2.end | jaccard | 50.3087224 | 0.68625834 | 0.001 | 0.00166667 | ** |
| Intake.water.2 vs SW.bioreactor.1 | jaccard | 18.0397039 | 0.60052869 | 0.001 | 0.00166667 | ** |
| Intake.water.2 vs Inoculum.1 | jaccard | 18.2686698 | 0.60355047 | 0.002 | 0.00264706 | ** |
| Intake.water.2 vs T1.start | jaccard | 21.9051885 | 0.48780974 | 0.001 | 0.00166667 | ** |
| Intake.water.2 vs T1.end | jaccard | 33.3104257 | 0.59154988 | 0.001 | 0.00166667 | ** |
| Intake.water.2 vs Intake.water.1 | jaccard | 12.1552087 | 0.50321274 | 0.001 | 0.00166667 | ** |
| Intake.water.2 vs Inoculum.2 | jaccard | 16.7482026 | 0.56299881 | 0.002 | 0.00264706 | ** |
| Intake.water.2 vs SW.bioreactor.2 | jaccard | 13.7808498 | 0.51457851 | 0.001 | 0.00166667 | ** |
| T2.end vs SW.bioreactor.1 | jaccard | 95.5153321 | 0.84890948 | 0.002 | 0.00264706 | ** |
| T2.end vs Inoculum.1 | jaccard | 64.702162 | 0.79192717 | 0.001 | 0.00166667 | ** |
| T2.end vs T1.start | jaccard | 48.0015353 | 0.63158639 | 0.001 | 0.00166667 | ** |
| T2.end vs T1.end | jaccard | 61.8119316 | 0.68823741 | 0.001 | 0.00166667 | ** |
| T2.end vs Intake.water.1 | jaccard | 55.3279097 | 0.76495933 | 0.001 | 0.00166667 | ** |
| T2.end vs Inoculum.2 | jaccard | 52.6011035 | 0.74504648 | 0.001 | 0.00166667 | ** |
| T2.end vs SW.bioreactor.2 | jaccard | 99.9751786 | 0.84742553 | 0.001 | 0.00166667 | ** |
| SW.bioreactor.1 vs Inoculum.1 | jaccard | 37.3395721 | 0.86155839 | 0.034 | 0.034 | * |
| SW.bioreactor.1 vs T1.start | jaccard | 11.7846413 | 0.40940727 | 0.001 | 0.00166667 | ** |
| SW.bioreactor.1 vs T1.end | jaccard | 37.2811288 | 0.68681565 | 0.001 | 0.00166667 | ** |
| SW.bioreactor.1 vs Intake.water.1 | jaccard | 26.6934098 | 0.81647677 | 0.03 | 0.03068182 | * |
| SW.bioreactor.1 vs Inoculum.2 | jaccard | 29.4575695 | 0.80799598 | 0.005 | 0.00642857 | ** |
| SW.bioreactor.1 vs SW.bioreactor.2 | jaccard | 50.3134884 | 0.8778647 | 0.008 | 0.00923077 | ** |

| | | | | | | |
|---|---|---|---|---|---|---|
| Inoculum.1 vs T1.start | jaccard | 2.50786434 | 0.12855658 | 0.026 | 0.0272093 | * |
| Inoculum.1 vs T1.end | jaccard | 23.4113226 | 0.57932582 | 0.001 | 0.00166667 | ** |
| Inoculum.1 vs Intake.water.1 | jaccard | 19.4518888 | 0.76426111 | 0.022 | 0.02357143 | * |
| Inoculum.1 vs Inoculum.2 | jaccard | 9.41562283 | 0.57357695 | 0.007 | 0.00851351 | ** |
| Inoculum.1 vs SW.bioreactor.2 | jaccard | 39.0270491 | 0.84791552 | 0.006 | 0.0075 | ** |
| T1.start vs T1.end | jaccard | 27.6372963 | 0.49674046 | 0.001 | 0.00166667 | ** |
| T1.start vs Intake.water.1 | jaccard | 11.5044258 | 0.40360139 | 0.001 | 0.00166667 | ** |
| T1.start vs Inoculum.2 | jaccard | 7.67242876 | 0.2988587 | 0.001 | 0.00166667 | ** |
| T1.start vs SW.bioreactor.2 | jaccard | 16.8805359 | 0.48395288 | 0.002 | 0.00264706 | ** |
| T1.end vs Intake.water.1 | jaccard | 14.2945634 | 0.45677466 | 0.002 | 0.00264706 | ** |
| T1.end vs Inoculum.2 | jaccard | 23.2201818 | 0.5633207 | 0.001 | 0.00166667 | ** |
| T1.end vs SW.bioreactor.2 | jaccard | 37.0801453 | 0.67320348 | 0.001 | 0.00166667 | ** |
| Intake.water.1 vs Inoculum.2 | jaccard | 16.698147 | 0.70461826 | 0.008 | 0.00923077 | ** |
| Intake.water.1 vs SW.bioreactor.2 | jaccard | 14.6650406 | 0.67689883 | 0.01 | 0.01097561 | * |
| Inoculum.2 vs SW.bioreactor.2 | jaccard | 29.5507754 | 0.78695513 | 0.01 | 0.01097561 | * |

**Supplementary Table S8.** Results of PERMANOVA pairwise comparison generated through 999 permutations between sample types and start- and end-time fraction of microeukaryotic (18S) communities in time series healthy and crash. Unweighted Jaccard was used as a distance metric. Sample type suffix denotes time series experiment: 1 = Time series crash, 2 = Time series healthy.

| Groups: 18S dataset | measure | F | R2 | p.value | p.adjusted | Significance |
|---|---|---|---|---|---|---|
| SW.bioreactor.1 vs Inoculum.1 | jaccard | 58.313788 | 0.90670741 | 0.03 | 0.03068182 | * |
| SW.bioreactor.1 vs T1.start | jaccard | 41.5047573 | 0.70942534 | 0.001 | 0.00155172 | ** |
| SW.bioreactor.1 vs T1.end | jaccard | 18.6966034 | 0.52376421 | 0.001 | 0.00155172 | ** |
| SW.bioreactor.1 vs Intake.water.1 | jaccard | 25.6345273 | 0.81033382 | 0.032 | 0.032 | * |
| SW.bioreactor.1 vs Inoculum.2 | jaccard | 53.6325077 | 0.88455038 | 0.008 | 0.009 | ** |
| SW.bioreactor.1 vs SW.bioreactor.2 | jaccard | 13.4141676 | 0.65710088 | 0.012 | 0.01285714 | * |
| SW.bioreactor.1 vs T2.start | jaccard | 41.4929616 | 0.70936674 | 0.001 | 0.00155172 | ** |
| SW.bioreactor.1 vs Intake.water.2 | jaccard | 19.7999977 | 0.62264148 | 0.003 | 0.00385714 | ** |
| SW.bioreactor.1 vs T2.end | jaccard | 96.1509094 | 0.84975817 | 0.001 | 0.00155172 | ** |
| Inoculum.1 vs T1.start | jaccard | 5.01010766 | 0.22762759 | 0.001 | 0.00155172 | ** |
| Inoculum.1 vs T1.end | jaccard | 15.0799108 | 0.47007334 | 0.001 | 0.00155172 | ** |
| Inoculum.1 vs Intake.water.1 | jaccard | 37.1256234 | 0.86087158 | 0.03 | 0.03068182 | * |
| Inoculum.1 vs Inoculum.2 | jaccard | 8.7607618 | 0.55585903 | 0.007 | 0.00828947 | ** |
| Inoculum.1 vs SW.bioreactor.2 | jaccard | 18.1982692 | 0.72220314 | 0.011 | 0.01207317 | * |
| Inoculum.1 vs T2.start | jaccard | 10.6996674 | 0.38627422 | 0.001 | 0.00155172 | ** |
| Inoculum.1 vs Intake.water.2 | jaccard | 21.8636655 | 0.64563789 | 0.003 | 0.00385714 | ** |
| Inoculum.1 vs T2.end | jaccard | 40.9654518 | 0.70672186 | 0.002 | 0.00272727 | ** |
| T1.start vs T1.end | jaccard | 43.9679736 | 0.61093805 | 0.001 | 0.00155172 | ** |
| T1.start vs Intake.water.1 | jaccard | 39.5134206 | 0.6991865 | 0.002 | 0.00272727 | ** |
| T1.start vs Inoculum.2 | jaccard | 9.98006442 | 0.35668483 | 0.001 | 0.00155172 | ** |
| T1.start vs SW.bioreactor.2 | jaccard | 34.9946565 | 0.66034311 | 0.001 | 0.00155172 | ** |
| T1.start vs T2.start | jaccard | 9.08627847 | 0.24500378 | 0.001 | 0.00155172 | ** |
| T1.start vs Intake.water.2 | jaccard | 51.6026021 | 0.69169976 | 0.001 | 0.00155172 | ** |
| T1.start vs T2.end | jaccard | 53.9636992 | 0.65838536 | 0.001 | 0.00155172 | ** |
| T1.end vs Intake.water.1 | jaccard | 13.0992315 | 0.43520153 | 0.001 | 0.00155172 | ** |
| T1.end vs Inoculum.2 | jaccard | 14.3909375 | 0.44428901 | 0.002 | 0.00272727 | ** |
| T1.end vs SW.bioreactor.2 | jaccard | 18.5838371 | 0.50797944 | 0.001 | 0.00155172 | ** |
| T1.end vs T2.start | jaccard | 43.1962267 | 0.60672073 | 0.001 | 0.00155172 | ** |
| T1.end vs Intake.water.2 | jaccard | 34.9700224 | 0.60324321 | 0.001 | 0.00155172 | ** |
| T1.end vs T2.end | jaccard | 46.9378455 | 0.62635702 | 0.001 | 0.00155172 | ** |

| | | | | | | |
|---|---|---|---|---|---|---|
| Intake.water.1 vs Inoculum.2 | jaccard | 27.1788463 | 0.79519496 | 0.008 | 0.009 | ** |
| Intake.water.1 vs SW.bioreactor.2 | jaccard | 11.9576476 | 0.63075587 | 0.007 | 0.00828947 | ** |
| Intake.water.1 vs T2.start | jaccard | 36.1444955 | 0.68011739 | 0.001 | 0.00155172 | ** |
| Intake.water.1 vs Intake.water.2 | jaccard | 26.9730782 | 0.69209514 | 0.002 | 0.00272727 | ** |
| Intake.water.1 vs T2.end | jaccard | 96.0079499 | 0.84956811 | 0.001 | 0.00155172 | ** |
| Inoculum.2 vs SW.bioreactor.2 | jaccard | 24.1070658 | 0.75083366 | 0.006 | 0.0075 | ** |
| Inoculum.2 vs T2.start | jaccard | 6.41699728 | 0.26280862 | 0.001 | 0.00155172 | ** |
| Inoculum.2 vs Intake.water.2 | jaccard | 28.6673292 | 0.68800496 | 0.001 | 0.00155172 | ** |
| Inoculum.2 vs T2.end | jaccard | 45.6985944 | 0.7174192 | 0.001 | 0.00155172 | ** |
| SW.bioreactor.2 vs T2.start | jaccard | 35.1160442 | 0.66111934 | 0.001 | 0.00155172 | ** |
| SW.bioreactor.2 vs Intake.water.2 | jaccard | 18.4585403 | 0.58675769 | 0.001 | 0.00155172 | ** |
| SW.bioreactor.2 vs T2.end | jaccard | 66.8063379 | 0.78775171 | 0.001 | 0.00155172 | ** |
| T2.start vs Intake.water.2 | jaccard | 51.351059 | 0.69065673 | 0.001 | 0.00155172 | ** |
| T2.start vs T2.end | jaccard | 46.6344592 | 0.62483817 | 0.001 | 0.00155172 | ** |
| Intake.water.2 vs T2.end | jaccard | 47.1667107 | 0.67220923 | 0.001 | 0.00155172 | ** |

**Supplementary Table S9.** Changes in prokaryotic communities between cultivation days within time series crash and healthy associated to total beta diversity (ßjac), turnover (ßjtu) and nestedness (ßjne) was inferred via PERMANOVA with 999 permutations using unweighted Jaccard distance.

| Dataset | Type | Df | SumOfSqs | MeanSqs | F.Model | R2 | P-value |
|---|---|---|---|---|---|---|---|
| **Crash/16S** | ßjac | 11 | 8.91380 | 0.81035 | 16.292 | 0.78874 | 0.001 |
| **Crash/16S** | ßjtu | 11 | 7.0898 | 0.64453 | 23.713 | 0.84458 | 0.001 |
| **Crash/16S** | ßjne | 11 | 0.19710 | 0.01792 | 2.3066 | 0.34581 | 0.035 |
| **Healthy/16S** | ßjac | 17 | 12.9770 | 0.76333 | 16.241 | 0.79316 | 0.001 |
| **Healthy/16S** | ßjtu | 17 | 10.8093 | 0.63584 | 27.886 | 0.86815 | 0.001 |
| **Healthy/16S** | ßjne | 17 | 0.0889 | 0.00523 | 0.60951 | 0.12581 | 0.765 |

**Supplementary Table S10.** Differences in measured bioreactor factors between Time series experiments: crash vs. healthy was tested using non-parametric Mann-Whitney U test.

| Factor | W-stat | p-value |
|---|---|---|
| Cells $L^{-1}$ | 122 | < 0.001 |
| Raw FL | 187 | < 0.001 |
| Bioreactor temperature ºC | 1214 | < 0.001 |
| O2 mg $L^{-1}$ | 150 | < 0.01 |
| pH level | 702.5 | 0.35 (ns) |
| DOC mg $L^{-1}$ | 204 | < 0.001 |
| $NO_2^-$ + $NO_3^-$ µM | 65 | < 0.001 |
| $PO_4^{3-}$ µM | 73.5 | < 0.01 |
| $Si(OH)_4$ µM | 55 | < 0.001 |

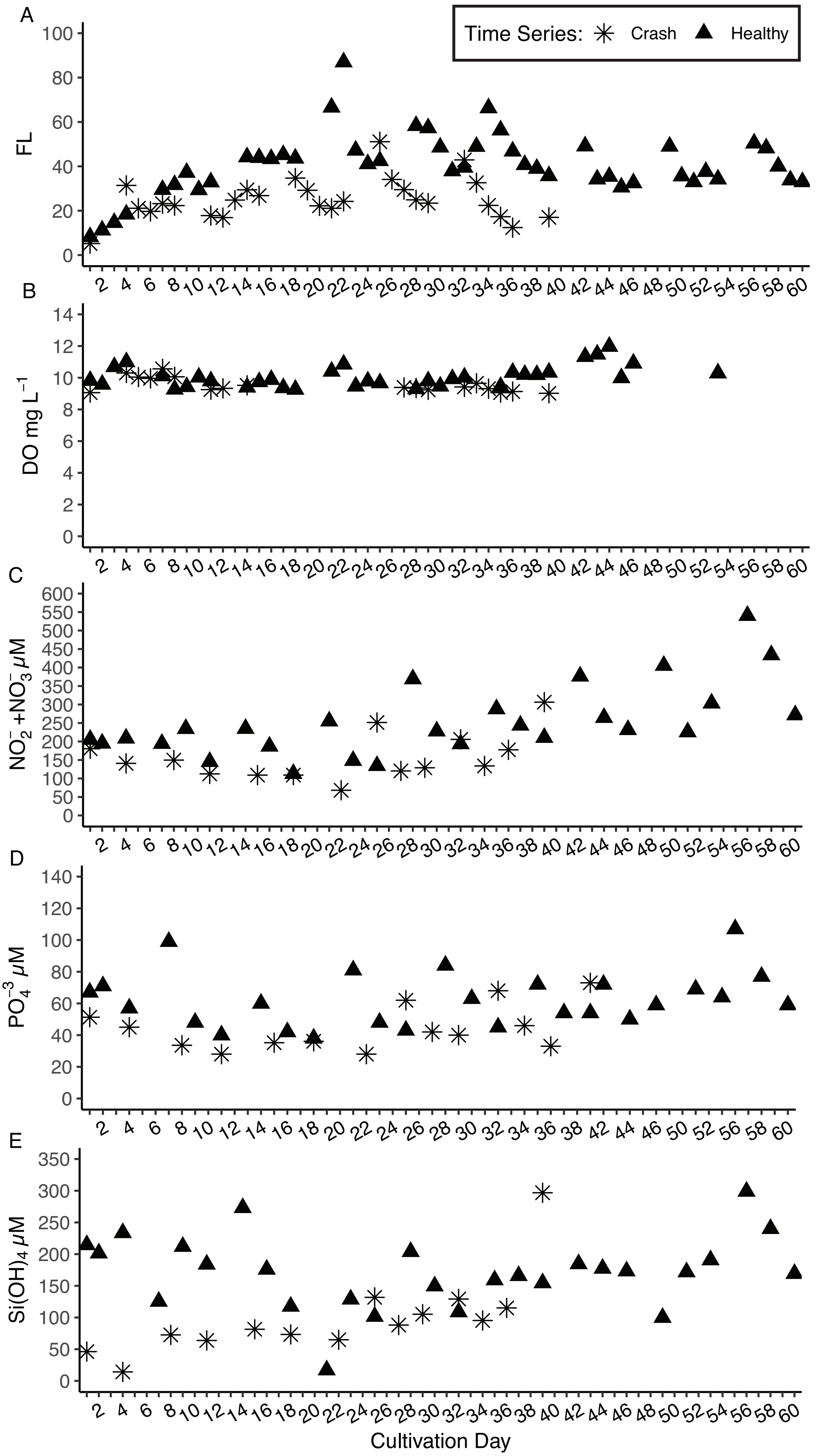


**Supplementary Figure S4.** Variation in bioreactor environment across time in crash and healthy cultivation. Daily or twice a week measurements of **(A)** raw fluorescence, **(B)** dissolved oxygen concentration, **(C)** combined nitrite and nitrate concentartion, **(D)** phosphate concentration and **(E)** silicate concentration. Note that the time series crash was ended after cultivation day 39.
